# How do mate-finding Allee effects affect evolutionary rescue?

**DOI:** 10.1101/2025.09.27.678992

**Authors:** Puneeth Deraje, Hildegard Uecker

## Abstract

Harsh environmental change can put populations at risk of extinction, requiring rapid adaptation for persistence. In sexually reproducing populations, the challenge of finding mates at low densities can impose a strong demographic Allee effect. When such a population falls below the Allee threshold, either through a reduction of the population size or an increase in the threshold, successful adaptation relies on overcoming the Allee effect, which gets increasingly difficult as the population declines. Despite mate-finding Allee effects being common, most models of evolutionary rescue assume that mating is assured even at low densities. Here, we set up a population genetic model for evolutionary rescue of a population below its Allee threshold. For the analysis, we combine stochastic computer simulations with mathematical arguments. As expected, mate limitation can severely impede rescue but the extent differs across sexual systems. We further show that mate limitation shifts the optimal sex ratio for dioecious populations, alters optimal evolutionary routes when there are trade-offs between increasing mate-finding efficiency and fecundity, and enhances the importance of standing genetic variation relative to *de novo* mutants. Overall, our results highlight the importance of accounting for positive density dependence in the assessment of a population’s scope for evolutionary rescue.

## Introduction

Populations exposed to a severe change in their environment are often maladapted to the new conditions and decline in size. Unless they rapidly adapt, they will eventually go extinct. This phenomenon of rapid adaptation that enables a population to survive a detrimental environmental change is termed evolutionary rescue (Gomulkiewicz and Holt, 1995). Studying evolutionary rescue is crucial for conservation, medicine, and agriculture for different reasons. For example, while evolutionary rescue is desirable from the conservation perspective (Carlson *et al*., 2014), the goal in medicine and agriculture is to prevent the escape of pathogens and pests from the effects of drugs and pesticides (Alexander *et al*., 2014; Kreiner *et al*., 2018).

While there now exists an extensive body of theory on evolutionary rescue, many of the facets of sexual reproduction and their role in rescue are still underexplored. Population genetic modeling studies account for intricacies like genetic linkage (Baskett and Gomulkiewicz, 2011; Uecker and Hermisson, 2016), sex-chromosome linked rescue (Unckless and Orr, 2020), and the mode of reproduction (Uecker, 2017). A common assumption in these (and other) models is that mating of individuals is assured. However, a key cost of sex is the need to invest into searching and choosing mates (Lehtonen *et al*., 2012). Shortage of mates can occur for a variety of reasons, such as low efficiency in searching for mates due to limited time or resources, a sparsely distributed population, short asynchronous reproductive cycles, limited mobility (for example, plants), reduction in pollinator populations (again in plants), and sperm limitation (Gascoigne *et al*., 2009). In ecological contexts, mate limitation has been extensively studied as a possible cause of Allee effects (positive correlation between individual fitness and population size) (Gascoigne *et al*., 2009; Berec, 2018; Boukal and Berec, 2009; Fauvergue, 2013; Shaw *et al*., 2018). Allee effects can also be of importance for eco-evolutionary dynamics, as demonstrated by Holt *et al*. (2004) who showed that Allee effects increase the importance of immigration for adaptation in a source-sink model. In the context of evolutionary rescue, mate limitation – the strength of which depends on both the abundance of mates and the efficiency of finding them – might be a strong obstacle to population survival. In order to survive, the population has to do both – adapt to the new environment and overcome the Allee effect.

Although the importance of Allee effects was already emphasized at the very inception of the term ‘evolutionary rescue’ (Gomulkiewicz and Holt, 1995), there are only few models of evolutionary rescue that incorporate Allee effects. These studies consider populations that are initially below the Allee threshold and ask when evolutionary rescue can occur via evolution of the threshold (Kanarek and Webb, 2010; Kanarek *et al*., 2015). While mate limitation as the cause for Allee effects is discussed by Kanarek and Webb (2010) and Kanarek *et al*. (2015), the Allee effect is modeled generically and the interest of these studies is the interaction with spatial processes rather than the interplay of mate limitation with relevant properties of sexually reproducing populations such as the sexual system and sex ratios.

By contrast, mate limitation is central in models of evolutionary rescue of plant populations that evolve higher selfing rates in response to pollen limitation (Cheptou, 2019; Xu, 2022, 2023). Since these studies assume that pollen limitation is due to a drop in pollinator density, pollen limitation is, however, independent of the population density. For example, Xu (2023) assumes that, independent of population size, a fixed proportion of ovules remains unfertilized. Density-dependent shortage of mates in a declining population could accentuate the benefits of selfing.

Another major assumption in evolutionary rescue models for sexually reproducing populations concerns the sexual system. Most models assume a hermaphroditic population (Kanarek *et al*., 2015; Uecker, 2017). However, a wide variety of sexual systems are known to exist in natural populations (The Tree of Sex Consortium, 2014). These include dioecious (two distinct sexes), androdioecious (males and hermaphrodites) (Pannell, 2002; Weeks, 2006), and gynodioecious (females and hermaphrodites) (Mc-Cauley and Bailey, 2009) populations. These sexual systems differ in the proportion of individuals producing male or female gametes, which could play a major role in rescue, especially in the presence of mate limitation. In the few studies that consider dioecious populations, the sex ratio is assumed to be 1:1 (Holt *et al*., 2004; Unckless and Orr, 2020). In other words, an offspring is female or male with equal probability. Although this assumption is true for many species and is explained by extensive theory (Fisher, 1958; Hamilton, 1967; Lloyd, 1974), biased sex ratios exist in natural populations (Barros *et al*., 2013; Székely *et al*., 2014). Again, when mating is not assured, the relative abundance of different sexes is expected to have a substantial effect on the probability of rescue.

In this study, we investigate the role of mate-finding Allee effects in evolutionary rescue. We assume that rescue is dependent on a one-locus 2-allele trait and set up an individual-based model motivated from branching processes with non-overlapping discrete generations. We apply this fundamental model to investigate the effects of a series of key factors that are expected to interact with the degree of mate-finding efficiency. Concretely, we provide: a comparison of rescue across dioecious, androdioecious, and hermaphroditic populations, an assessment of the importance of standing genetic variation vs *de novo* mutations, a comparison of rescue via increased efficiency in finding mates vs increased fecundity, an analyis of the effect of the sex ratios in dioecious and androdioecious populations, and a brief exploration of the effect of selfing in androdioecious populations with varying degrees of inbreeding depression. Overall, the results demonstrate that mate limitation has crucial effects on the dynamics of evolutionary rescue.

### The Model

We consider a population of sexually reproducing haploid individuals with discrete non-overlapping generations. The population has been subjected to a change in its environment and is declining in size. However, it can be rescued from extinction by a mutation occurring at a single locus. We thus have two genotypes – the wild type and a mutant. The mutant may or may not pre-exist prior to the environmental change.

We consider three sexual systems: (1) dioecy, where individuals are either male or female, i.e. they produce either male or female gametes, (2) hermaphroditism, where each individual produces both male and female gametes, and (3) androdioecy, where the population is composed of hermaphrodites and males. As summarized in Table 1, populations of the three sexual systems differ in the sizes of their reproductive pool (individuals producing female gametes) and mate pool (individuals producing male gametes).

**Table 1:**
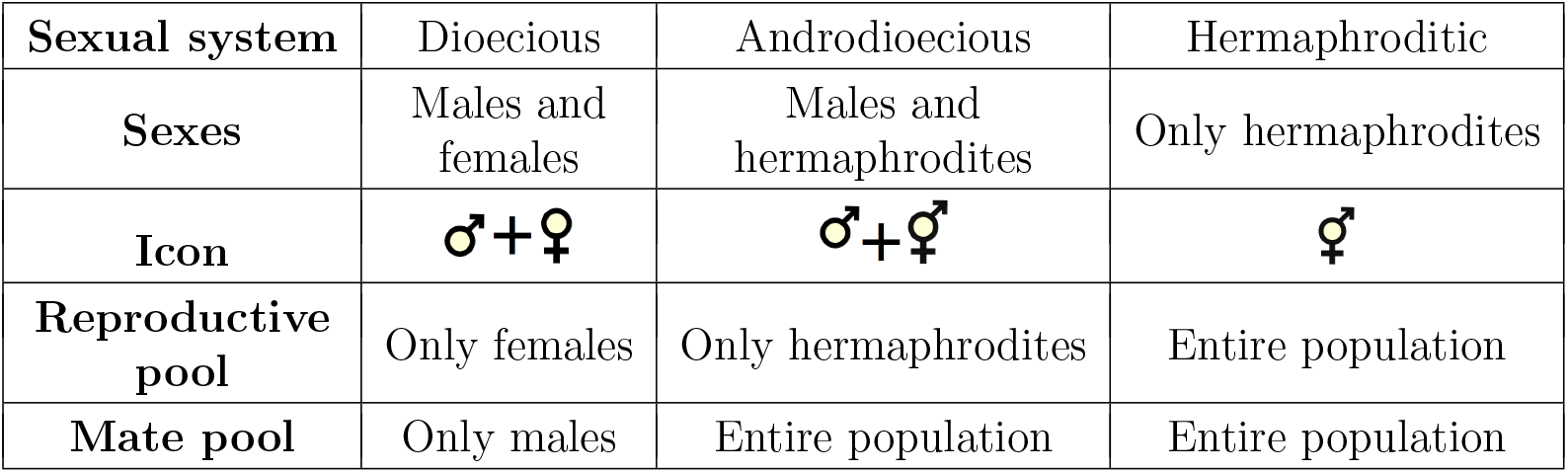
Overview of the three sexual systems.

In all three cases, the life cycle is modeled by the following scheme that extends the branching process for a sexually reproducing population as described by Daley (1968) to include a density-dependent mate-finding probability and two genotypes (wild type and mutant). In the following, we often speak of females and males, which for hermaphrodites should be understood as “individuals in the female or male role”. In generation *t*, each female (or hermaphrodite in the female role) in the population encounters at least one potential mate with a mate-finding probability 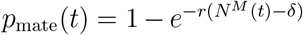, where *N*^*M*^ (*t*) is the total number of individuals with male gametes in the population and the parameter *r* quantifies the efficiency of finding mates; *δ* = 0 for dioecious populations and *δ* = 1 for hermaphroditic and androdioecious populations as we exclude selfing. For dioecious populations, the form of this mate-finding probability can be derived by assuming that the probability of encountering a mate while searching an infinitesimal extra area is proportional to that area and is either independent or linearly dependent on the number of previous encounters (Dennis, 1989; Boukal and Berec, 2002; Berec, 2018). An underlying assumption is furthermore that a male can mate with arbitrarily many females. We extend this approach to hermaphroditic individuals by ignoring that an encounter between two hermaphrodites is a simultaneous mate finding success for two females, i.e. we assume that the mate encounter probabilities are independent from each other; formally, this is correct if we assume that females search sequentially rather than simultaneously. For any sexual system, while a female might encounter more than one mate, we assume that she only mates once. Which male she encounters and mates with is random. Consequently, the probability that the mating male will be of a given genotype is proportional to the frequency of males of that genotype. Upon mating, the number of offspring produced by a mating pair is distributed according to a Poisson distribution with mean *λ*_*w*_ or *λ*_*m*_ (with *λ*_*m*_ *> λ*_*w*_), depending on whether the mother is a wild-type or mutant individual respectively. The offspring inherits the genotype of either of its parents with equal probability. Further, a wild-type offspring can acquire the mutation with probability *µ*. In a dioecious population, the offspring is a female with probability *α*_fem_ and male otherwise. In an androdioecious population, the offspring is hermaphroditic with probability *α*_her_. A diagram of the life cycle is shown in Fig. 1 for the dioecious population and in Fig. S1 for the two other sexual systems.

**Figure 1:**
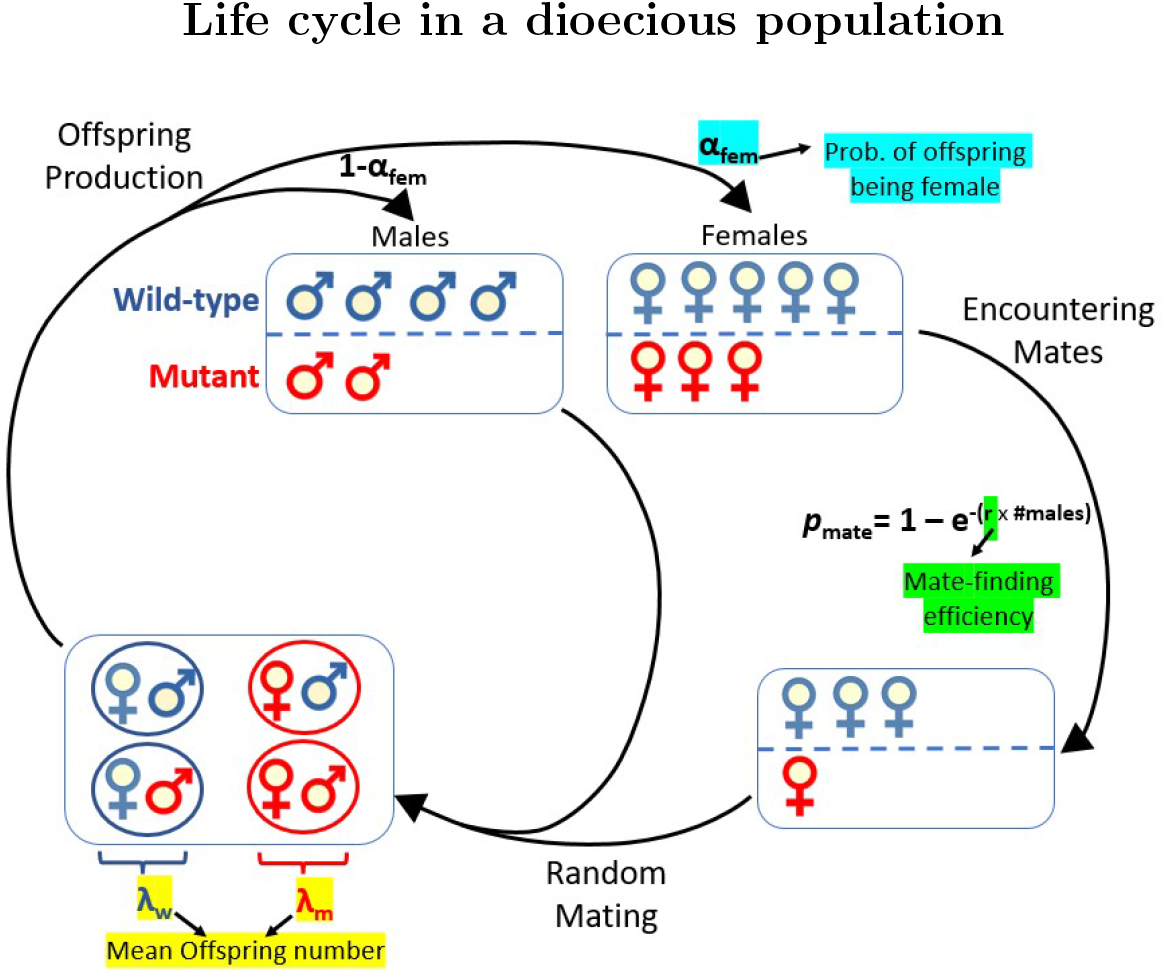
Diagrammatic representation of the stochastic life cycle of a dioecious population to go from generation *t* to *t* + 1 (excluding selfing and mutation). The blue individuals have the wild-type phenotype and the red ones have the mutant pheno-type.

We furthermore consider two extensions of the model. First, in the version described above, which we will apply for most of the manuscript, we assume that the mutant and the wild type only differ in their fecundity. However, the benefit of the mutant could also – at least partially – stem from an increased efficiency in searching for mates, and we will compare both routes to rescue below. Second, since hermaphroditic individuals have both male and female reproductive organs, there may also be a possibility of selfing in androdioecious and hermaphroditic populations, and we briefly explore the consequences of selfing at the end of the Results section.

The formal descriptions of the models for all three populations are given in the SI section S1.

### Analysis and Results

The model described in the previous section was implemented using the open source software *Python* 3.8. For computing the rescue probability, unless stated otherwise, we start with a population of *N*_0_ = 10^4^ individuals with *α*_fem_*N*_0_ females for a dioecious population and *α*_bis_*N*_0_ hermaphrodites for an androdioecious population. We run simulations until either the population goes extinct or the mutation occurs and the total population reaches a size of 2*N*_0_ or 2 · 10^4^, whichever is larger. Technically if the population hits *N*_0_ post-decline its a good indication that the population survived but in some simulations (especially with low initial population size or low decline rate), the population can hit *N*_0_ immediately after the change but go extinct later. Therefore, to avoid such scenarios, we chose the cutoff for successful rescue as 2*N*_0_ or 2 · 10^4^. We always perform 10^5^ replicate simulations; the fraction of populations that survive give the rescue probability.

### Characterizing the regimes for population decline and evolutionary rescue

Prior to exploring the factors affecting evolutionary rescue, we quantify the regime in which the wild-type population is maladapted such that population decline is certain. This regime is characterized by the critical wild-type fecundity 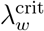 below which the wild-type population has a zero probability of survival (in the absence of any mutants). For a hermaphroditic population in which mating is assured it holds that 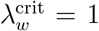, since in that case the dynamics can be approximated by a simple branching process for which the critical value is one (Harris, 1963). Similarly, for dioecious and androdioecious populations, given guaranteed matings, the critical values are 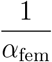 and 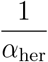 (see Theorem 1 in Daley (1968) for a proof). We derive an approximation for the critical value in the presence of mate limitation under the assumption that the initial number of individuals producing female gametes is *α*_rep_*N*_0_, where *α*_rep_ represents the proportion of offspring that belong to the reproductive pool. To do so, we determine the criteria for a decrease in the expected population size in the first generation after the environmental change; if *λ*_*w*_ is too small to maintain the population size in the first generation, it is also too small in all later generations. We obtain for the expected number of wild-type individuals in generation *t* = 1

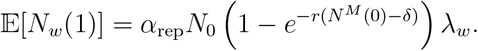

It holds that 𝔼 [*N*_*w*_ (1)] *< N*_0_ if 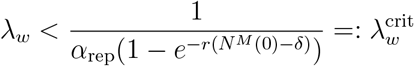. Specifically,

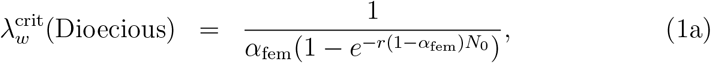

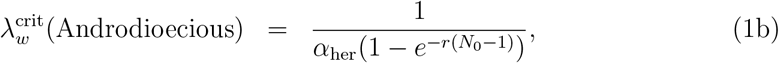

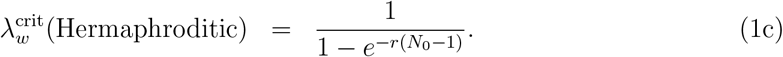

To verify these approximations with simulations, we determine the survival probability for a fully wild-type population in the absence of mutation (*µ* = 0). For this, we increase the fitness of wild-type individuals *λ*_*w*_ in steps of 0.01, starting from zero, until we encounter a non-zero survival probability; the corresponding wild-type fecundity is the critical value 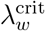. A comparison between approximation (1) and simulation results shows that the approximations work well for large initial population sizes but overestimate 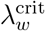 for small populations, in which demographic stochasticity cannot be neglected (Fig. 2a and b).

**Figure 2:**
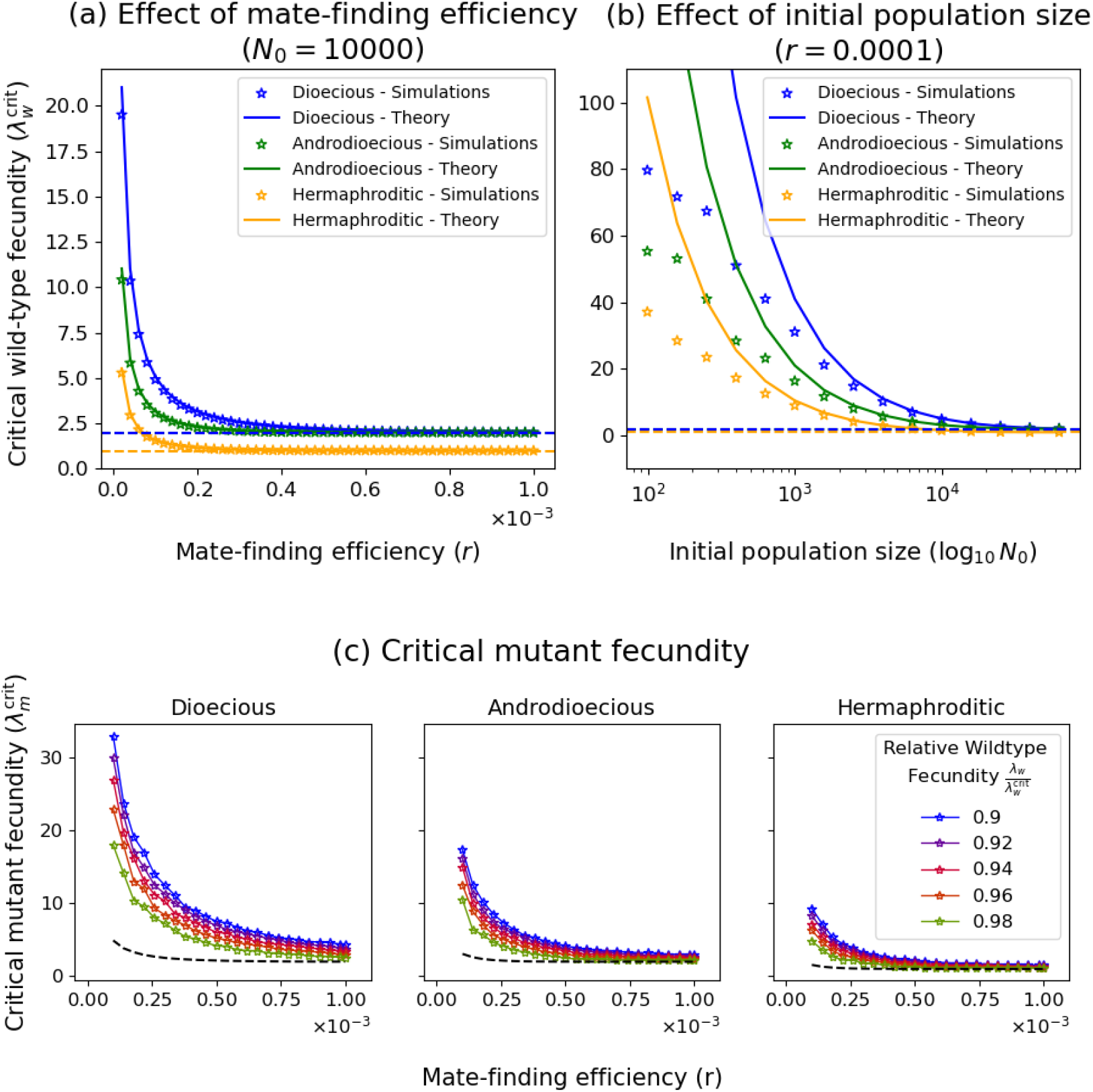
Critical wild-type and mutant fecunditities. The upper row illustrates the dependence of the critical wild-type fecundity 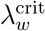 on (a) the mate-finding efficiency *r* and (b) the initial population size *N*_0_ for the three sexual systems. The solid lines are the deterministic approximation (Eq. 1) and stars show results from stochastic simulations as described in the main text. For 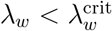, the population goes extinct with certainty in the absence of evolution. In the limit of assured mating (*r* → ∞ or *N*_0_ → ∞), the critical wild-type fecundity 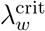 converges to 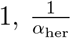, and 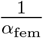 for the hermaphroditic, the androdioecious, and the dioecious populations respectively. Panel (c) shows the minimum mutant fecundity required for a non-zero rescue probability 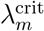 as a function of the mate-finding efficiency (*r*) for different scaled wild-type fecundities 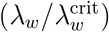. The dashed black line is the critical wild-type fecundity as a function of the mate-finding efficiency and is included for comparison. Simulation points are connected by lines to guide the eye. Other parameter values: *N*_*w*_(0) = 10^4^, *N*_*m*_(0) = 0, *α*_fem_ = 0.5, *α*_her_ = 0.5. For (a)-(b) *µ* = 0 and (c) *µ* = 10^−4^.

In the limits *r* → ∞ or *N*_0_ → ∞, approximation (1) recovers the known results for 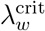 for populations with assured mating. In this limit, the critical values of the dioecious and androdioecious populations eventually converge to the same value if *α*_fem_ = *α*_her_. In contrast, in the presence of mate limitation, the androdioecious population has a lower critical value, since the mate pool is larger (Fig. 2a and b). Finally, the hermaphroditic population always has the lowest critical value since it has the largest mate and reproductive pools of the three populations. In brief, 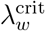 is a function of the mate-finding efficiency *r*, the initial population size *N*_0_, the sexual system, and the sex ratio. For finite *r*, as we will explain in more detail in the Discussion section, a population with 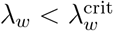 is below its Allee threshold.

Further, note that the sexual system will affect the rate at which a wild-type population decays for the same wild-type fecundity (Fig. S2a). If the wild-type fecundity scaled by the critical value 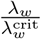 is the same, decay rates are similar across sexual systems (Fig. S2b).

After having characterized the fecundity range in which the wild-type population is maladapted, we also explored the minimum mutant fecundity required to observe a non-zero rescue probability if rescue relies on *de novo* mutations (i.e. in the absence of standing genetic variation). This critical mutant fecundity 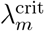 was computed in a similar fashion as the critical wild-type fecundity 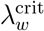: We computed the rescue probability for a given wild-type fitness *λ*_*w*_ and mate-finding efficiency *r*, increasing the mutant fitness in increments of 0.01 (starting from zero) until we encountered a non-zero rescue probability. The corresponding value of *λ*_*m*_ is the critical mutant fecundity 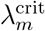. At low mate-finding efficiencies the critical mutant fecundity is much higher than the critical wild-type fecundity, indicating the severe consequences of mate limitation on survival of populations (Fig. 2c and S4).

### Preliminary considerations

The probability of evolutionary rescue depends on both the availability of rescue mutants and their probability to successfully spread and not get lost while rare. Rescue mutants might either already pre-exist prior to change (rescue from the standing genetic variation 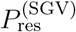) or arise *de novo* during population decline (rescue from *de novo* mutations 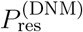). The total probability of evolutionary rescue is given by

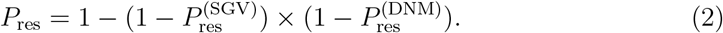

Following the classic population genetics approach to determining probabilities of rescue (e.g. Orr and Unckless, 2008), we can write

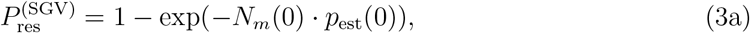

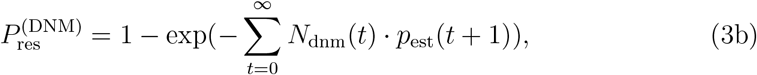

where *N*_*m*_(0) is the number of mutants at time *t* = 0 and 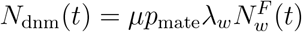 is the number of new mutations that get generated from generation *t* to *t*+1 (where *p*_mate_ depends on the number of wild-type males in the population). The establishment probability *p*_est_(*t*) is the probability that a mutant present at time *t* establishes a long-term lineage of offspring that rescues the population; time dependence occurs here through dependence on the declining wild-type population size (i.e. small *N*_*w*_ corresponds to large *t*). The underlying assumption of this approach is that mutant lineages suffer independent fates. If mutants occur so rarely that mating between different mutant lineages can be ignored during the establishment phase, this is a good assumption. Mutations might appear in either females/hermaphrodites or males in dioecious/androdioecious populations, and *p*_est_ in Eq. (3) is the weighted average.

We will not calculate *N*_dnm_(*t*) and *p*_est_(*t*) directly. Instead, we choose a computationally simpler approach to obtain insights into how model parameters affect the rate of appearance and spread of mutants. By considering the expected changes in the number of wild-type individuals from one generation to the next, Δ*N*_*w*_, in the absence of mutants we can learn how fast the wild-type population decays in dependence of model parameters and thus how those parameters affect the rate of appearance of new mutants. We further use the expected changes in the number of mutants, Δ*N*_*m*_, as a proxy to understand effects on their establishment (establishment is of course also affected by the variance in offspring numbers, which we ignore here). Given the simplicity of the expressions for Δ*N*_*w*_ and Δ*N*_*m*_, this approach proves very helpful in explaining – and predicting – simulation results. We here provide the expressions for Δ*N*_*w*_ (before the appearance of mutants) and Δ*N*_*m*_ (omitting the expectation and time dependence for notational simplicity) and refer to the Supplementary Information section S2.1 for details. For a dioecious population, we obtain

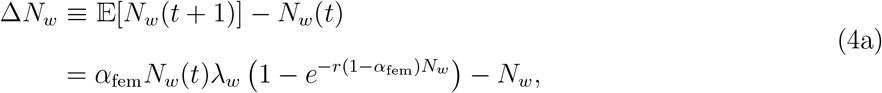

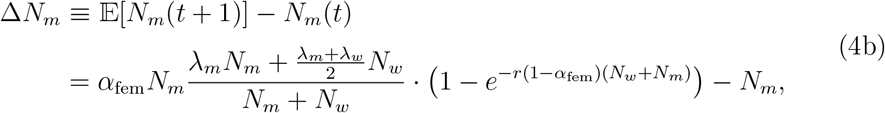

where we assume that the proportions of females and males are given by *α*_fem_ and 1 − *α*_fem_. For an androdioecious population, we have

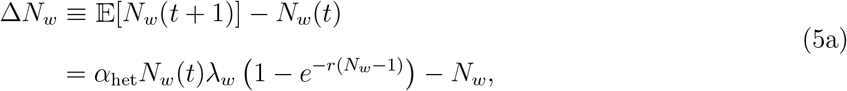

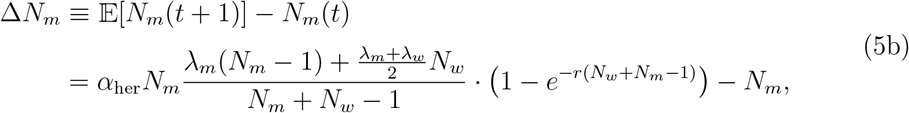

which also holds for a hermaphroditic population with *α*_her_ = 1.

As a side remark, we point out that for a dioecious population, the deterministic frequency increase of the mutant allele is the same as that of a beneficial allele with selection coefficient *s* = (*λ*_*m*_ − *λ*_*w*_)*/λ*_*w*_ and a dominance coefficient 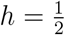 in a diploid randomly mating population (see section S2.2).

### Comparison of the probability of rescue by *de novo* mutations across the three sexual systems

We first explore the probability of rescue by *de novo* mutations for the three sexual systems across a broad parameter range (Fig. 3). Similar to the findings for the critical wild-type fecundity 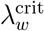, we observe that hermaphroditic populations have the highest rescue probability followed by androdioecious populations, and dioecious populations have the lowest chances of survival. As expected, for all three systems, the rescue probability decreases with the strength of mate limitation, albeit to different extents. The relative advantage that a hermaphroditic population has over a dioecious population is higher for a low (*r* = 0.0001) than for a moderate mate-finding efficiency (*r* = 0.001). The reason is that a hermaphroditic population has a larger mate pool and a larger reproductive pool, which both provide a benefit when the mate finding probability is low. In contrast, when the mate finding probability is high, the benefit of a larger mate pool is negligible such that the advantage over the dioecious population is less pronounced. Besides these qualitative insights, Fig. 3 also allows us to obtain a quantitative account of the severe constraints that mate limitation can exert on rescue and demonstrates that very high mutant fecundities might be required for the population to have a decent chance to survive. This is of course particularly true if the wild-type decays quickly and the mate-finding efficiency is low, and is furthermore particularly the case for a dioecious population. Nevertheless, for moderate mate limitation and wild-type fecundities close to the critical wild-type fecundity, rescue is possible even with moderate mutant fecundities (see especially insets in Panel a).

**Figure 3:**
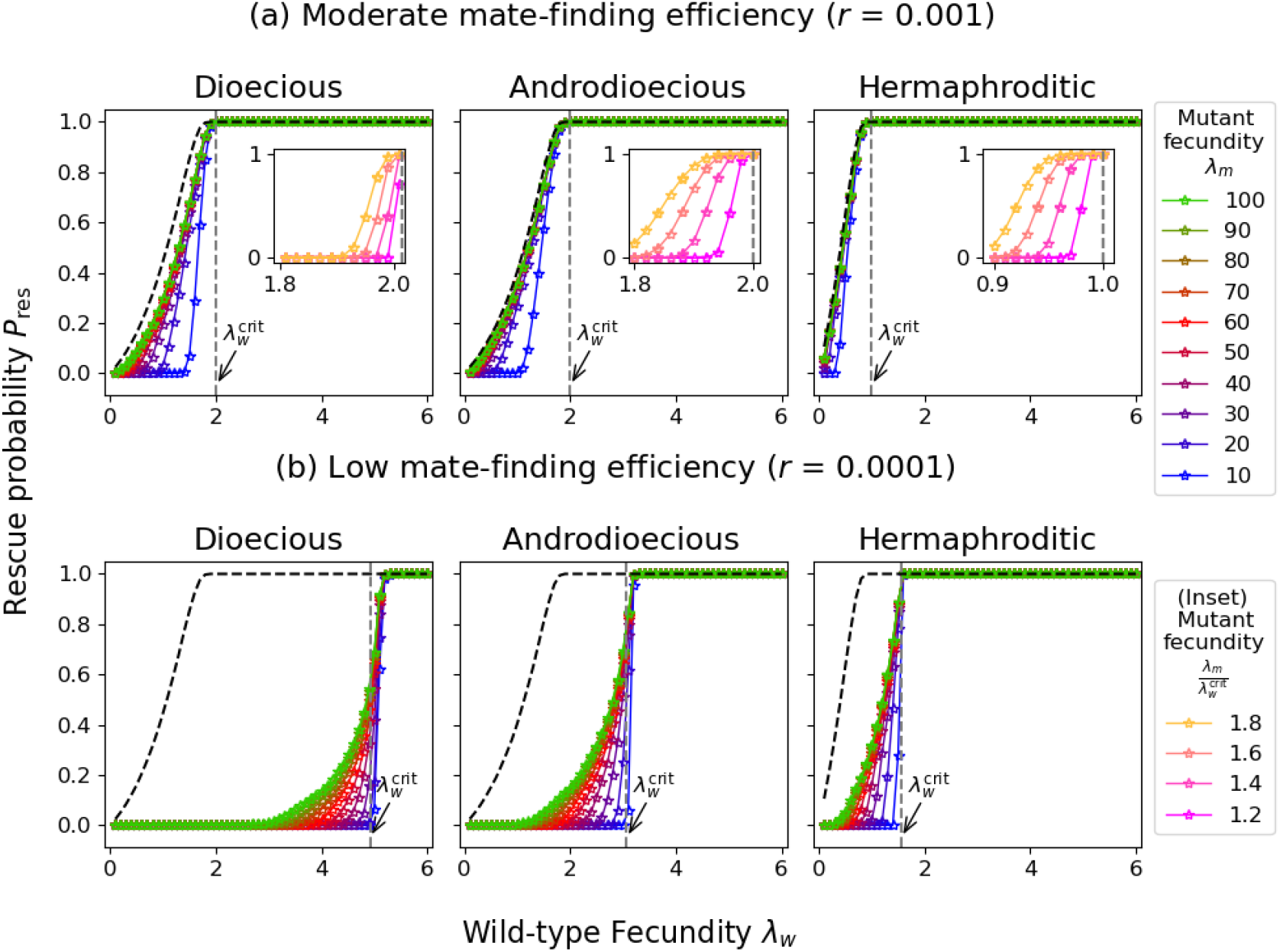
Rescue probability by *de novo* mutations for dioecious, androdioecious, and hermaphroditic populations for a moderate (*r* = 0.001) and a low (*r* = 0.0001) mate-finding efficiency. The dashed black lines show the rescue probability when mating is assured for a mutant fecundity of *λ*_*m*_ = 10 (curves for higher *λ*_*m*_ coincide since for large *λ*_*m*_, any mutant that appears will establish if mating is guaranteed; rescue thus only depends on the probability of appearance of mutants, which is independent of *λ*_*m*_). The vertical dashed line indicates the critical wild-type fecundity 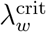, which depends on the mate-finding efficiency and has been obtained from stochastic simulations (see Fig. 2). A lower mate-finding efficiency leads to a lower rescue probability for all three sexual systems. The rescue probability is highest for the hermaphroditic population owing to both high mate and reproductive pools. The insets in Panel a zoom into a range of wild-type fecundities between 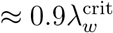 and 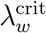, where rescue is likely even for a moderate mutant fecundity. Simulation points are connected by lines to guide the eye. Other parameters: *N*_*w*_(0) = 10^4^, *N*_*m*_(0) = 0, *µ* = 10^−4^, for the androdioecious population *α*_her_ = 0.5, for the dioecious population *α*_fem_ = 0.5.

### Rescue due to standing genetic variation versus recurrent mutation

In Fig. 3, rescue relies on *de novo* mutations. However, the rescue mutant might already pre-exist in the population prior to environmental change. In Fig. 4, we compare for a dioecious population the probability of rescue by a single mutant pre-existing in the standing genetic variation 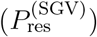 with that of rescue by recurrent *de novo* mutations 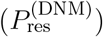. As expected, rescue from new mutations is more likely than rescue from the standing genetic variation if the wild-type decays slowly (i.e. if the wild-type fecundity is close to the critical wild-type fecundity 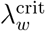) but is smaller otherwise (for a very low mate-finding efficiency as in Panel c, rescue from the standing genetic variation is more likely than *de novo* rescue across the entire simulated range). Perhaps less obvious, we further observe that the wild-type fecundity 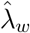 for which 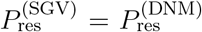 depends on the mutant fecundity, and it does so in opposite ways with and without mate limitation: For assured mating, the dependence is weak with a slight shift of 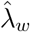 to lower values for decreasing mutant fecundities (Panel a). By contrast, 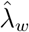 substantially shifts to higher values with increasing *λ*_*m*_ if mates are limited (Panel b).

**Figure 4:**
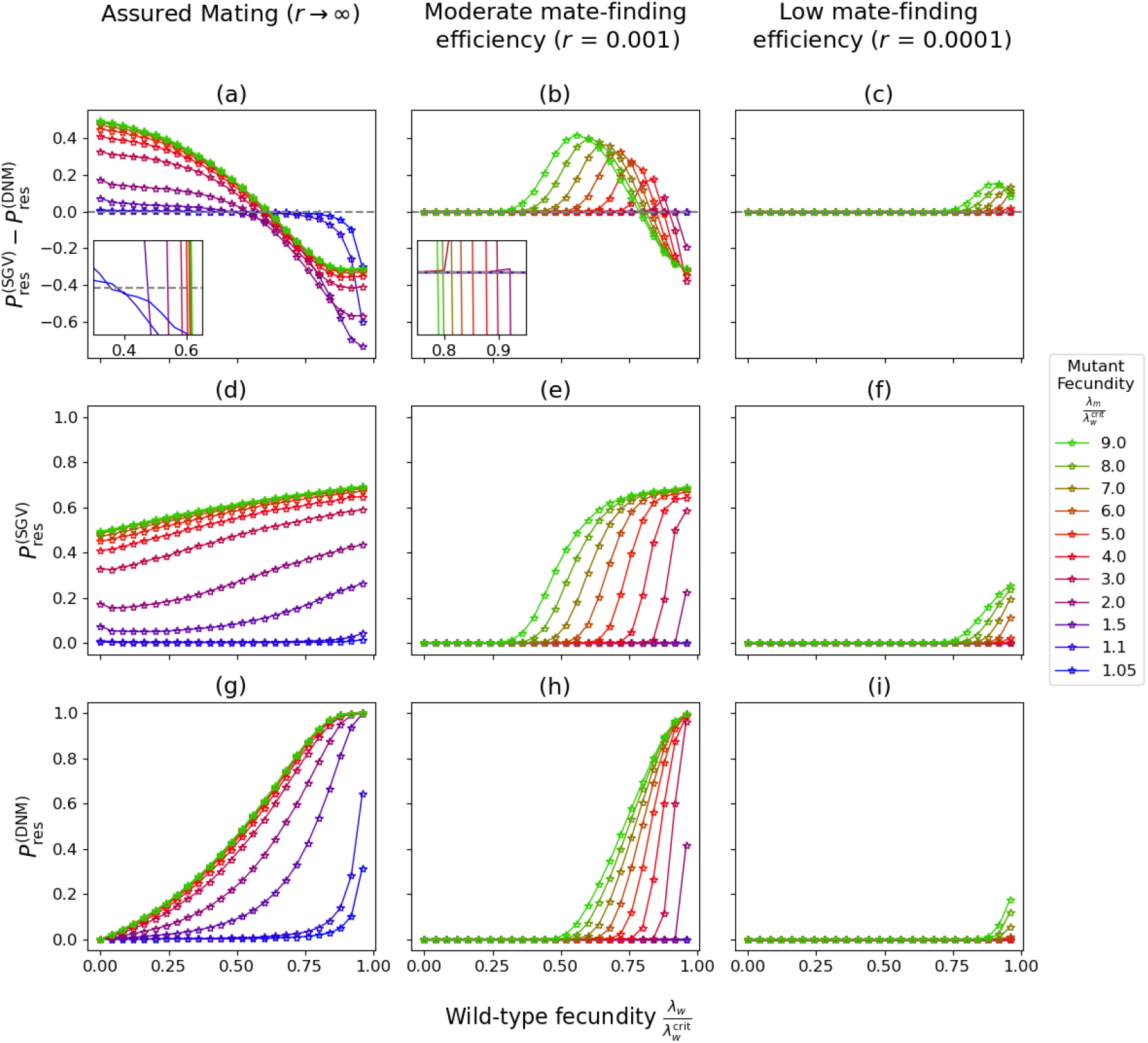
Comparison of the probabilities of rescue from the standing genetic variation 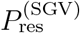 and from *de novo* mutations 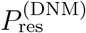 in a dioecious population. The top row shows the difference between the two probabilities, while the middle and bottom rows show the absolute probabilities. The columns show results for assured mating (left column), a moderate mate-finding efficiency (middle column), and a low mate-finding efficiency (right column). A lower mate-finding efficiency leads to a larger range of (scaled) wild-type fecunditites *λ*_*w*_ in which rescue from the standing genetic variation is more likely than rescue from *de novo* mutations. Further, the range in which 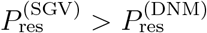 depends on the mutant fecundity *λ*. To determine 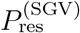, we set *N*_*m*_(0) = 1 (with the mutant individual being female with probability *α*_fem_) and *µ* = 0; to determine 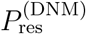, we set *N*_*m*_ (0) = 0 and *µ* = 10^−4^. Other parameter: *α*_fem_ = 0.5, *N*_0_ = 10^4^.

To understand this pattern, it is insightful to think of the problem in terms of Eq. (3). If the establishment probability *p*_est_ was independent of the wild-type population size at the time of appearance, the two rescue probabilities would be equal when the number of new mutations equals that of mutants in the standing genetic variation, independent of *λ*_*m*_. By contrast, if the establishment probability depends on the wild-type population size, the crossing point 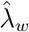 shifts away from this point. More precisely, if the establishment probability decreases with the wild-type population size, early mutants enjoy an advantage over late occurring mutants and the intersection of 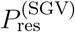 and 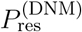 shifts to the right, and vice versa. Thus, how does *p*_est_ depend on the wild-type population size for assured and for limited mating? We here provide an intuitive explanation with further details given in Supplementary Material sections S2.3 and S2.4.

If mating is assured, mutants are able to reproduce independent of the wild-type population size (given there is a mutant to mate with). However, they are more likely to mate with each other when *N*_*w*_ is small. A mutant female mating with a wild-type male has *λ*_*m*_*/*2 mutant offspring, and a mutant male mating with a wild-type female has *λ*_*w*_*/*2 mutant offspring. If they mate with each other, however, they have together *λ*_*m*_ *>* (*λ*_*m*_ + *λ*_*w*_)*/*2 mutant offspring. For this reason, Δ*N*_*m*_ as given in Eq. (4b) decreases with increasing wild-type population size. This effect is not very relevant if *λ*_*m*_ is large as in that case any mutant female will have many offspring anyway. However, mating of mutants with each other boosts establishment when *λ*_*m*_ is small, and the establishment probability therefore decreases with time as the wild-type population size drops (see also Fig. S5). Therefore, we see a shift of the crossing point to the left for small *λ*_*m*_. The same effect occurs in principle with mate limitation. However, there is a much stronger negative effect of a decreasing wild-type population size on *p*_est_: With mate limitation, the spread of the mutant crucially depends on the ability to find mates, and the larger the wild-type population size by the time the mutation occurs, the more likely it is to do so (Δ*N*_*m*_ in Eq. (4b) decreases with increasing *N*_*w*_.) Consequently, mutants that already pre-exist prior to the environmental change profit from the initially large population size, and the wild-type fecundity 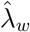 at which 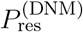 equals 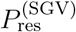 shifts to the right. Again, it shifts further for smaller *λ*_*m*_. This is because once a female mutant has reproduced, the (expected) number of mutants is larger with larger *λ*_*m*_, and further establishment therefore depends less on the mating success per mutant; moreover, in the extreme of very large *λ*_*m*_, mating might be assured through the presence of sufficiently many mutants. Overall, the range of wild-type fecundities where rescue by standing genetic variation is more likely than rescue by new mutations is larger for limited than for assured mating.

Finally, we observe in Fig. 4 that not only 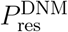 but also 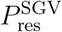 increases with *λ*_*w*_, irrespective of the mate-finding efficiency. While a rapidly disappearing wild type and thus low *λ*_*w*_ promotes advantageous matings between mutants (see above), a high wild-type fecundity is beneficial in any one mating that occurs between a wild-type female and a mutant male, i.e. large *λ*_*w*_ has a direct positive effect on Δ*N*_*m*_. A very rapid elimination of the wild-type (and thus very low *λ*_*w*_) can be beneficial if the initial number of mutants is sufficiently large that they do not rely on matings with wild-type individuals in their early phase of spread: if we set *N*_*m*_(0) = 10 instead of *N*_*m*_(0) = 1, we observe a minimum in 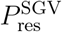 for some small *λ*_*w*_, which appears as a result of the two conflicting effects of *λ*_*w*_ (see Fig. S6). Otherwise, the same general conclusions hold for *N*_*m*_(0) = 10 as for *N*_*m*_(0) = 1.

### Comparing the effect of increased fecundity versus higher mate-finding efficiency on the probability of rescue

So far, we have assumed that the mutant has a higher fecundity than the wild type but that its mate-finding efficiency *r*_*m*_ is the same (i.e. *r*_*m*_ = *r*_*w*_ = *r*). However, a mutant can also rescue the population from extinction if it is more efficient at finding mates, even if its fecundity is unaltered. Therefore, we now allow the mutant to differ from the wild type in both parameters, considering a dioecious population (Fig. 5). Specifically, we focus on a scenario where individuals have limited resources (like energy and time) to invest. Consequently, there is a trade-off between searching for food (where we consider higher food uptake as a proxy for higher fecundity) or searching for mates, and the changes in *λ*_*m*_ and *r*_*m*_ invoked by a mutation of a given effect size are constrained by this trade-off. Let the mutant fecundity and mate-finding efficiency be given by *λ*_*m*_ = (1 + *s*_*λ*_)*λ*_*w*_ with *s*_*λ*_ ≥ 0 and *r*_*m*_ = (1 + *s*_*r*_)*r*_*w*_ with *s*_*r*_ ≥ 0. The trade-off is then modeled by the assumption *s*_*λ*_ + *s*_*r*_ = *c* for some constant *c*.

**Figure 5:**
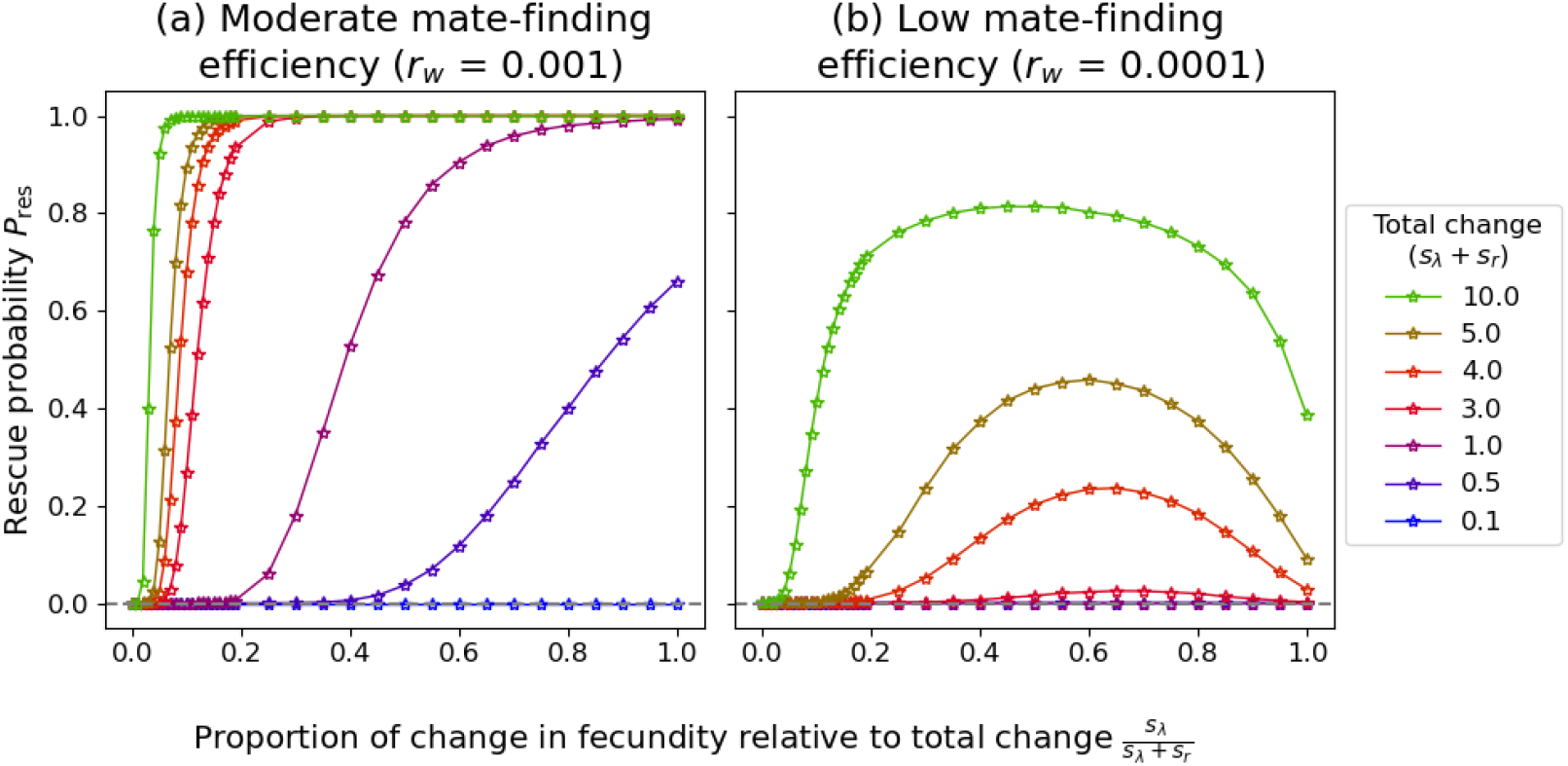
Rescue probability of a dioecious population when the mutant differs from the wild type in both its fecundity and its mate-finding ability. Increases in fecundity and mate-finding efficiency are assumed to be subject to a trade-off, and the panels show the rescue probabilities as a function of the proportion of the mutational effect that is allocated to an increase in the fecundity. The lines show results for different total effect sizes of the mutation. If the wild-type mate-finding efficiency *r*_*w*_ is large, a mutant with increased fecundity is more likely to rescue the population than one with increased mate-finding efficiency (Panel a). In contrast, if the wild-type mate-finding efficiency *r*_*w*_ is low, a mutant that increases both quantities to some extent is most favorable (Panel b). Simulation points are connected by lines to guide the eye. Other parameters: 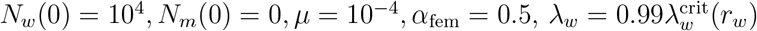.

We find that when the wild-type mate-finding efficiency *r*_*w*_ is high, rescue is more likely by a mutation that increases fecundity than by one that increases mate finding, i.e. the probability of evolutionary rescue *P*_res_ increases with *s*_*λ*_ for *s*_*λ*_+*s*_*r*_ = *c* (Fig. 5a). However, when the wild-type mate-finding efficiency *r*_*w*_ is low, the rescue probability is highest for intermediate values of *s*_*λ*_, as improving the ability to find mates is also crucial (Fig. 5b).

### The influence of the sex ratio on rescue

In sexual systems, like androdioecy and dioecy, where individuals differ in whether they produce male or female gametes or both, the sex ratio is expected to be a key determinant of a population’s ability to survive harsh changes as it determines the sizes of the reproductive and mate pools. We therefore proceed to investigate the effect of the sex ratio on rescue probabilities in the interplay with mate limitation. We vary the sex ratio in the dioecious system by changing the (expected) proportion of females (*α*_fem_) and in an androdioecious system by varying the (expected) proportion of hermaphrodites (*α*_her_). Note that in a dioecious system the sex ratio – individuals producing female gametes divided by individuals producing male gametes – will be 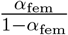, while for an androdioecious system it is given by *α*_her_ since all individuals produce male gametes. Therefore, the sex ratio in androdioecious systems can never exceed one (while it can be female-biased in dioecious populations).

We first consider a dioecious population (Fig. 6 a-c). When mating is assured, the rescue probability is highest for *α*_fem_ close to one (Panel a). However, the lower the mate-finding efficiency, the further the optimal sex ratio shifts towards a 1:1 ratio, i.e. *α*_fem_ = 0.5 (Fig. 6 top row from left to right). Intuitively, the reason for this shift in the optimal sex ratio is that with lower mate-finding efficiency, it becomes more important to have larger numbers of males so that females are able to find them and mate. In contrast, when mating is assured, having more females directly contributes to higher offspring numbers and therefore female-biased ratios are favoured (the fraction of females, *α*_fem_, cannot be exactly one since there always needs to be at last one male for mating). We can understand the reasons behind the respective optimal sex ratios in more detail by following the approach outlined in the section “Preliminary considerations”. This in particular means that we decompose the problem into the rate at which new mutants appear, *N*_dnm_(*t*), and their establishment probability, *p*_est_(*t*). We draft the argument here; detailed calculations can be found in the SI section S2.5.

**Figure 6:**
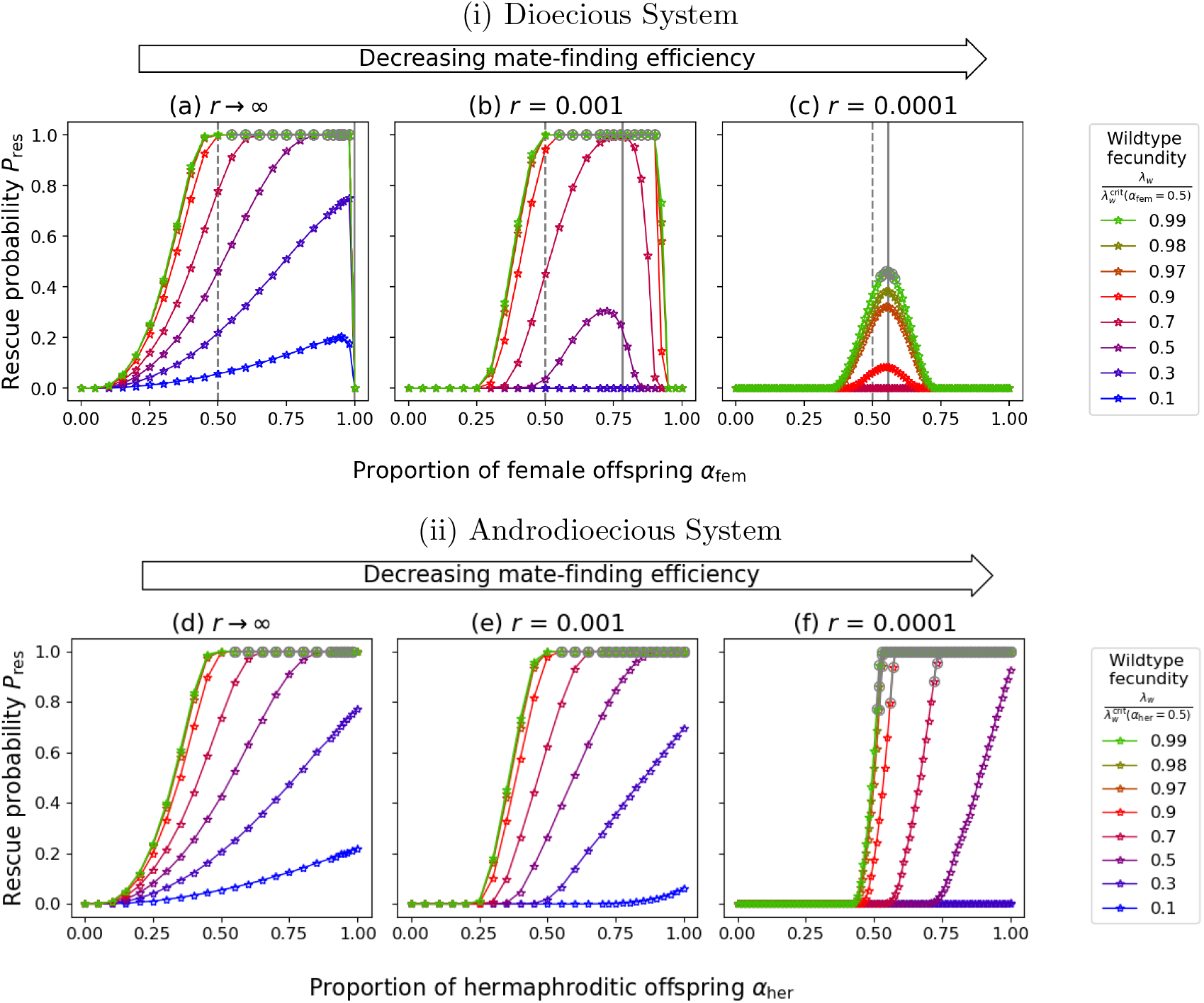
Dependence of the rescue probability on (i) the proportion of females *α*_fem_ in a dioecious population and (ii) the proportion of hermaphrodites *α*_her_ in an andro-dioecious population for different degrees of mate-finding efficiency (a-c and d-f). The grey solid line for the dioecious system (a-c) is the value of the sex ratio that minimizes the critical wild-type fecundity given in Eq. 1a and the dashed grey line indicates a balanced sex ratio for comparison. The grey circles are points for which the wildtype is above the critical wild-type fitness. Simulation points are connected by lines to guide the eye. Other parameters: 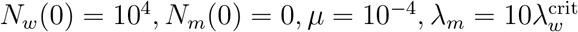.

New mutations appear at rate

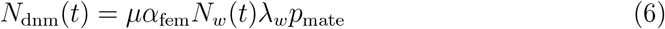

With 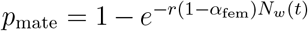. For assured mating, *p*_mate_ = 1 (provided there exists at least one male); for a low mate-finding efficiency *r*, Taylor expansion in *r* leads to the approximation *p*_mate_ ≈ *r*(1 − *α*_fem_)*N*_*w*_(*t*). Thus

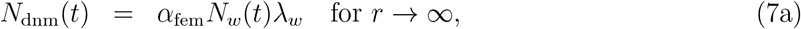

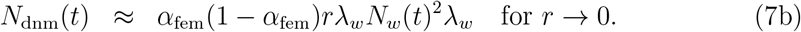

The direct dependence of *N*_dnm_(*t*) on *α*_fem_ is thus consistent with the observed optimal sex ratio in Fig. 6. We further note that the wild-type population size *N*_*w*_ also decays slowest for *α*_fem_ = 1 if mating is assured and for *α*_fem_ = 1*/*2 in the limit *r* → 0 (see Eq. 4a). (Accordingly, the critical wild-type fecundity, Eq. (1), is lowest for *α*_fem_ = 1 if mating is assured and for *α*_fem_ = 0.5 if mate limitation is severe.) The fraction of females that maximizes the rate of appearance of new mutations is thus 1 and 1*/*2 for assured mating and severe mate limitation, respectively. As a proxy for the dependence of the establishment probability on *α*_fem_, we consider the expected change in the number of mutants from one generation to the next (Eq. 4b). With the same approximation of *p*_mate_ as before, we see that the dependence of Δ*N*_*m*_ on *α*_fem_ is again in line with the optimal values observed in Fig. 6. The indirect dependence on *α*_fem_ via the wild-type population size is somewhat more subtle. A slow decay of the wild-type population favors spread of the mutant for low *r*. Therefore, an even sex ratio (*α*_fem_ = 1*/*2) is consistently optimal across the direct and indirect effects on *p*_est_. By contrast, for assured mating (*r* → ∞), we have seen in sections S2.4 and S2.3 that a low wild-type population size can be beneficial for spread of the mutant by enabling mating between mutant females and males. Since the wild-type decays faster for lower *α*_fem_, it is conceivable that the establishment probability can be highest for *α*_fem_ *<* 1 if mating of mutants is essential for their establishment. If we consider such a regime (*λ*_*w*_ and *λ*_*m*_ rather small), we indeed observe that the peak in the rescue probability is at somewhat lower sex ratios (Fig. S7a).

For an androdioecious population, irrespective of the mate-finding efficiency *r*, the rescue probability is always highest for *α*_her_ = 1, which is a population of only hermaphrodites (Fig. 6 d-f). This is because hermaphrodites contribute to both the reproductive and the mate pool. A formal argument as for the dioecious population can be found in the SI section S2.5.

In brief, in both sexual systems, mate limitation favours an even sex ratio (Fig. 6c and f). However, the manner in which this is achieved differs. In a dioecious system, an even sex ratio occurs when the proportion of female individuals is 1*/*2 (Fig. 6c), whereas in an androdioecious system, it is achieved if all individuals are hermaphroditic (Fig. 6f). When finding mates is assured, a bias towards the production of female gametes maximizes the probability of rescue. In a dioecious population, this implies that a female-biased population (*α*_fem_ ⪅ 1) has the highest chance of survival. For androdioecy, *α*_her_ = 1 does not only lead to an even sex ratio, but simultaneously leads to the largest possible number of individuals producing female gametes.

### The effect of selfing and inbreeding on rescue

We finally investigate the extent to which selfing can compensate for shortage of mates in an androdioecious population. We thereby assume that hermaphroditic individuals that are unable to find mates self with probability *p*_self_. We account for inbreeding depression due to selfing in a simplistic manner by multiplying the mean number of offspring of a selfing individual by a factor (1 − ID_self_). The results (Fig. 7) indicate that the option to self leads to a substantial increase in the rescue probability. Even for strong inbreeding depression, selfing still somewhat increases the rescue probability because even for high ID_self_ the selfing individuals still have offspring, which is better than none. We observe that as expected, the effect of selfing is particularly pronounced when finding mates is difficult (low *r*) and inbreeding depression low (small ID_self_). A population with a low mate-finding efficiency and a high probability to self can have a higher rescue probability than a population with a higher mate-finding efficiency but lower selfing probability.

**Figure 7:**
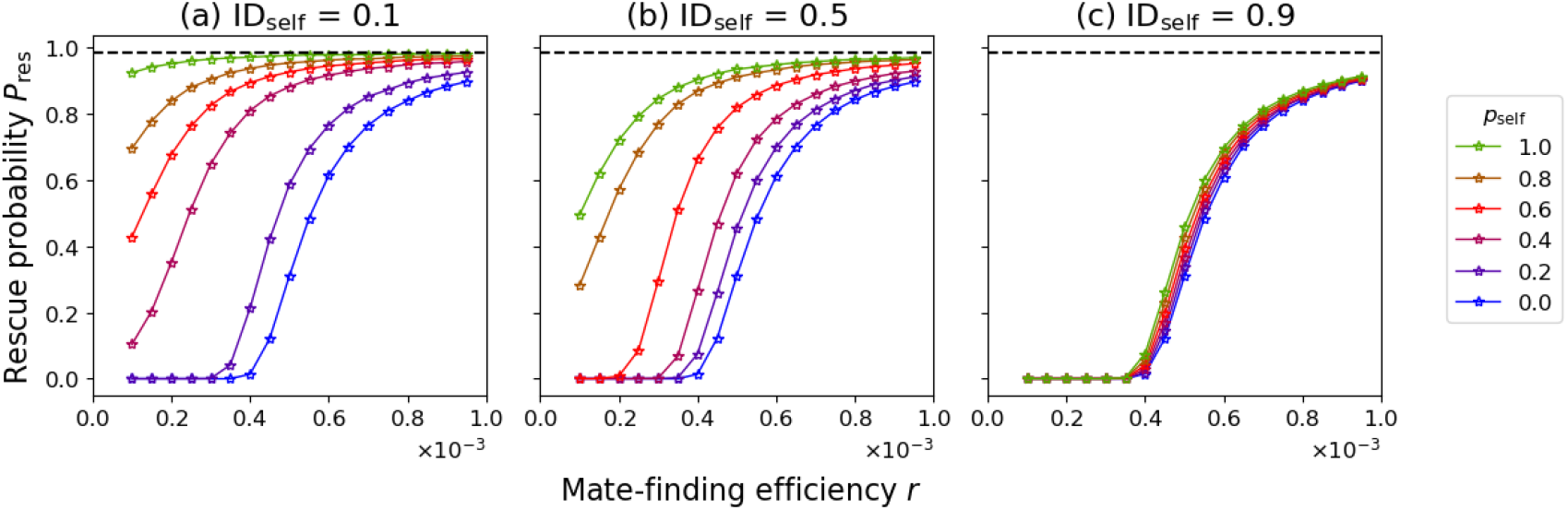
Rescue probability as a function of the mate-finding efficiency for different probabilities of selfing (colors of solid lines) and increasing degree of inbreeding depression (left to right) in an androdioecious population. The dashed black line is the rescue probability for a population with assured mating for the same wild-type and mutant fecundity. The rescue probability increases with increasing probability of selfing and decreasing inbreeding depression. Other parameters: 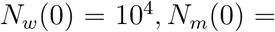 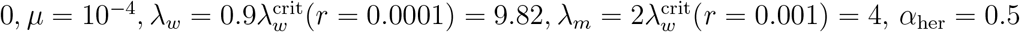.

## Discussion

Harsh environmental change can cause a population to fall below its Allee threshold, either through a one-time event, gradual population decline, or an increase of the Allee threshold beyond the current population size. A particularly important Allee effect for sexually reproducing populations might appear through density-dependent mate limitation. In this article, we set up an evolutionary rescue model to determine the probability that a population that finds itself below its mate-finding Allee threshold is able to escape extinction through adaptive evolution. Since both the reproductive and the mate pools matter for population growth, the constraints that mate limitation imposes on rescue differ between sexual systems and are, for a given total population size, particularly strong for dioecious populations. As the strength of mate limitation, modulated by the mate-finding efficiency, increases, the relative importance of the two pools shifts, leading to a shift in the optimal sex ratio in dioecious populations. Similarly, the relative importance of adaptation via increased fecundity vs increased mate-finding efficiency changes with the extent of mate limitation. As overcoming the Allee effect becomes more and more difficult as the population size decreases, standing genetic variation is particularly relevant for rescue in populations experiencing an Allee effect.

### The Allee threshold

Mate limitation leads in our model to a component Allee effect, i.e. a positive relationship between population size/density and some component of fitness (Stephens *et al*., 1999), which is here the expected number of offspring. Since we do not have any other density-dependent effect (such as increased offspring survival at low densitites) which could compensate for the mate-finding Allee effect, the population also experiences a demographic Allee effect, i.e. a positive relationship between population density and population growth (Stephens *et al*., 1999; Courchamp *et al*., 2008). More precisely, we model a strong Allee effect, i.e., the population growth rate is negative below a threshold population size (Kramer *et al*., 2009). Below this Allee threshold, population extinction is certain in the absence of evolution. In our model, the Allee threshold of a monomorphic population (dropping subscripts on the parameters) is given by

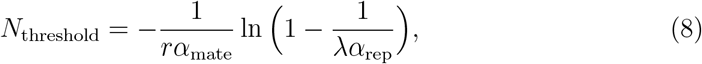

which is for a given sexual system dependent on the intrinsic fecundity, the sex ratio, and the mate-finding efficiency. Consequently, environmental change affecting any of these three components could increase the threshold such that the current population falls below the new threshold, leading to population decline towards extinction. Similarly, any of the three life-history parameters could evolve to rescue the population from extinction. For most of our analysis, we focus on the evolution of higher fecundity as a pathway to rescue.

Non-specific Allee effects are often modeled by making the population growth rate – typically in a Lotka-Volterra framework – proportional to a factor (*N/a* − 1). In such models, the Allee threshold *a* is – unlike in our model – an independent model parameter. This is the formulation assumed by Kanarek and Webb (2010) and Kanarek *et al*. (2015), who study evolutionary rescue via evolution of the Allee threshold in a spatial context. Here, the value of *a* evolves without affecting the maximal population growth rate or any other life-history parameter.

Generally, it is expected that populations have evolved to avoid mate-finding Allee effects (Rankin and Kokko, 2007; Gascoigne *et al*., 2009). Despite the interesting questions that this raises, the evolution of Allee thresholds (irrespective of cause) has rarely been modeled even outside a rescue context. An exception is Berec (2018) who study how the mate search rate and the Allee threshold evolve under various scenarios. Importantly, they find that under many circumstances, evolution at higher population densities leads to a higher Allee threshold than evolution at lower densities. This suggests, as they point out, that anthropogenically small populations may have through their evolutionary history at large densities – trait values that make them susceptible to extinction once rare. This hypothesis underscores the importance of accounting for mate-finding Allee effects in evolutionary rescue studies.

### The relative importance of standing genetic variation vs *de novo* mutants

Once the population is declining, it is not sufficient to restore population fecundity to the critical wild-type fecundity (see the high critical mutant fecundities in Fig. 2c). A mutant can only be successful if the current population size is above its Allee threshold - indeed, it needs to be well above it since most individuals in the population are still unfit wild-type individuals and this wild-type population size keeps declining. There is thus a limited window of opportunity for evolution to occur. This has consequences for the relative importance of standing genetic variation vs *de novo* mutations.

Orr and Unckless (2014) characterized the condition under which rescue from the standing genetic variation is more likely than rescue by *de novo* mutations in an asexually reproducing haploid population in which the establishment of the rescue mutant is independent of the wild-type population size. This condition is fulfilled when the number of mutants in the standing genetic variation is larger than the expected number of new mutants during population decline (Eq. 4 in Orr and Unckless, 2014). In our scenario, the mating probability decreases with the population size and early mutants therefore have a much higher chance to spread than late mutants. Consequently, the importance of standing genetic variation for surviving harsh environmental changes is particularly high in populations displaying mate-finding Allee effects. Alternatively, immigration from a source could maintain a wild-type population in the sink, facilitating establishment of a mutant population. Such a scenario has been analysed in Holt *et al*. (2004).

Perhaps more surprisingly, even if mating is assured, the wild-type population size has some effect on establishment of mutants in sexually reproducing populations – albeit a weak one in comparison: a mutant female and a mutant male leave more mutant offspring together than when each of them mates with a wild-type partner. Yet, pairing of mutants is more likely at low wild-type population sizes. Therefore, by contrast to a population with a mate-finding Allee effect, where the negative impact of a low population size dominates, establishment of late mutants is more likely than that of early ones if mating is assured. A similar negative effect of the wild type on the spread of the rescue genotype has previously been highlighted in haploid 2-locus and diploid 1-locus models (Uecker and Hermisson, 2016; Uecker, 2017). In those scenarios, mating between the rescue genotype – the double mutant or the mutant homozygote – and the wild type breaks the rescue type up such that only a quarter of the offspring are again rescue mutants.

### Trade-offs between fecundity versus mate searching

For most of our study, we assume that rescue occurs via evolution of increased fecundity. However, an increase in the mate-finding efficiency lowers the Allee threshold as well and could lead to rescue. To test which trait should best evolve, we set up a scenario in which both are allowed to evolve but account for a trade-off between them. Examples of such trade-offs can be found in natural populations. For instance, the ability to take flight and thus find more mates is known to trade off with reproductive output in multiple insects (Langellotto *et al*., 2000; Chang *et al*., 2021). When resources, like energy or time, are limited, foraging effort needs to be split between finding mates or finding food, with the latter impacting fecundity (Prokopy and Roitberg, 1984; Blanckenhorn *et al*., 1995). In our rescue model, we consider a mutation with an unconditionally beneficial effect: While we limit the increase in fecundity and mate-finding efficiency by the trade-off, the mutant is at least as good in both traits as the wild type, and both traits can simultaneously improve to some extent. Incorporating this trade-off, we find that adaptation via increased fecundity is better than adaptation via increased mate-finding efficiency when mate limitation is weak anyway. However, when mate limitation is severe, an intermediate increase in both fecundity and mate finding is ideal (Fig. 5).

Asking about the evolution of mate-search rates and Allee thresholds, Berec *et al*. (2018) set up a model in which an increase in the mate-search rate is accompanied by a decrease in fecundity. They find that irrespective of a trade-off, the mate-search rate always evolves to an intermediate value when females are the searching sex, as in our model. However, when males are the searching sex, mate searching evolves to ever higher values, and in the presence of a trade-off with fecundity, the population ultimately goes extinct (an instance of evolutionary suicide). Adapting our model to males being the searching sex would be an interesting future avenue.

### The role of the sexual system and the sex ratio

Sexual systems differ in the sizes of their mate and reproductive pools. One of the major advantages of hermaphroditism over dioecy is thought to be the higher mate assurance when there is low availability of mates, either due to low densities or due to low mobility (Pannell and Jordan, 2022). Consistent with this, we find that the rescue probability drops much more severely with decreasing mate-finding efficiency for dioecious than for hermaphroditic populations (Fig. 3). In our model, we assume that hermaphrodites do not face a trade-off between investing into female and male functions. It has been speculated that such trade-offs could be absent or weak when both functions benefit from the same investment or rely on different resources (Pannell and Jordan, 2022). In other cases, they might be substantial, which would diminish the benefit of hermaphroditism over the other two sexual systems. We furthermore find that critical fecundities as well as rescue probabilities can be similar for different sexual systems when mates are assured but start to differ when mates are harder to find, as seen by the results for dioecious and androdioecious populations (see Fig. 2 and 3).

Besides the sexual system, populations may also differ in their sex ratios. Explaining observed sex ratios by evolutionary principles has been the focus of theoretical studies for a long time. To explain the commonly observed 1:1 sex ratio in dioecious populations, Fisher (1958) proposed that in populations with biased sex ratios the rarer sex would have a higher fitness and therefore would become more common such that the evolutionary stable sex ratio would be 1:1. While later work crucially extended sex ratio theory (Hamilton, 1967; Bull and Charnov, 1988, e.g.), the number of studies taking mate limitation into account is limited (Lehtonen and Schwanz, 2018). Lehtonen and Schwanz (2018) showed that under panmixia, a balanced sex ratio (1:1) is always the evolutionarily stable strategy, irrespective of mate limitation. While Lehtonen and Schwanz (2018) focus on the evolution of sex allocation, our model assesses the ability of a population to survive an extinction-level event, given a fixed operational sex ratio. In that case, the optimal sex ratio depends on the mate-finding efficiency and thus the degree of mate limitation at low densities. When mate limitation is severe, a balanced sex ratio minimizes the risk that a populations is wiped out by harsh environmental change, while for assured mating, female-biased populations have an advantage. While not the most prominent driver of sex ratios, the respective rates of population extinction may contribute to the distribution of sex ratios seen in nature.

### Future directions

Our model is, of course, based on many assumptions, both concerning the ecology and the genetics. First, our model assumes a specific mating system (unlimited polygyny). Extending it to other mating systems such as monogamy or polyandry or to a limited the number of matings per male will likely yield new results. For example, it has been found that the sex ratio that maximizes the female mating rate depends on the mating system (Bessa-Gomes *et al*., 2004). Further complexities such as female choosiness or a need of multiple matings to fertilize all eggs alter the Allee effect as well with likely consequences for the probability of evolutionary rescue. While incorporating these aspects individually in evolutionary rescue models is interesting in itself, developing a general framework based on an effective degree of mate limitation (which would need to be determined for each scenario) would be a major advancement.

We further consider a very simple life cycle. More complex life cycles with additional effects of density such as additional component Allee effects (Berec *et al*., 2007) or conversely increased survival of offspring at low densities could reveal interesting and unexpected behaviors. Similarly, males and females could have different death rates throughout the mating season, for example because they are differentially affected by environmental change such as males being hunted more, which would lead to changing sex ratios over time (Boukal and Berec, 2009).

On the genetic side, we have considered the most simple genetic basis – a mutation at a single locus. Considering polygenic adaptation is an important next step. Furthermore, inbreeding is an important consideration, which we have only touched upon in a very simplified manner. Decreases in population size due to mate-finding Allee effects can enhance inbreeding, causing a genetic Allee effect, cascading into an extinction vortex (Wittmann *et al*., 2018). A detailed model of inbreeding in diploid individuals could be applied to assess how much such effects reduce the probability of rescue and how the advantages and disadvantages of selfing balance out.

Last, incorporating the possibility of evolutionary rescue into population dynamics models with Allee effects has important applications. Allee effects have received much attention by modelers in the context of conservation, assessing extinction times of small populations (e.g. Bessa-Gomes *et al*., 2004) or – in few studies – the effect of warming temperatures on Allee thresholds in specific systems (Wittmann *et al*., 2011; Berec, 2019). Yet, these studies are purely ecological and do not take the evolutionary potential of populations into account. Similarly, models for pest-control strategies that exploit Allee effects ignore possible escape from extinction through evolution (Boukal and Berec, 2009; Tobin *et al*., 2011). Our results show that it is difficult but not impossible for populations below their Allee threshold to adapt and recover. Evolutionary models could help to design more effective conservation programs that take the evolutionary potential of populations into account – or alternatively, to prevent invasion of species whose Allee threshold has been lowered through environmental change. They could especially provide insights into which control strategies that introduce or enhance an Allee effect make escape of invasive specise, agricultural pests, or disease vectors from interventions unlikely. For example, introduction of sterile males “only” requires adaptations to overcome the mate-finding Allee effect as in our model. On the other hand, driving a population below its mate-finding Allee threshold through pesticide application requires two adaptations – one to overcome the Allee effect and the other to become resistant to the pesticide – but only under continued application of pesticides.

## Conclusion

To summarize, we developed a general theoretical model to analyse evolutionary rescue of a population that fell below its mate-finding Allee threshold. We provided a quantitative assessment of the limits to evolutionary rescue imposed by mate limitation, which depend on the sexual system, and analyzed how mate limitation interacts with a range of relevant factors. Our results show that mate limitation can alter the relative importance of mate pool versus reproductive pool, adaptation via standing genetic variation versus *de novo* mutation, and adaptation via increased fecundity versus higher mate-finding efficiency. These results have implications for fundamental questions in evolutionary biology such as those on the observed diversity of sexual systems and sex ratios as well as for conservation. We hope that our work builds a starting point for future studies, including models tailored for specific purposes such as contributing to species risk assessment or designing pest control strategies.

## Acknowledgments

The authors thank Pete Czuppon, Kuangyi Xu, Christin Nyhoegen, and the Osmond Lab for helpful discussions. The simulations were performed on the Computer cluster at the Max Planck Institute for Evolutionary Biology and on the Niagara supercomputer at the SciNet HPC Consortium. SciNet is funded by: the Canada Foundation for Innovation; the Government of Ontario; Ontario Research Fund – Research Excellence; and the University of Toronto.

