## Supplemental file for "How do mate-finding Allee effects affect evolutionary rescue?"

### S1 Detailed model description

#### S1.1 Dioecious population

Let $N_j^i(t)$ $(i = M, F\ ,\ j = w, m)$ denote the number of individuals in generation $t$ of type $j$ ($w$ild-type or $m$utant) and gender $i$ ($M$ale or $F$emale). Further, $N_j(t) = N_j^M(t) + N_j^F(t)$ will denote the total number of individuals of type $j$ and $N^i(t) = N_w^i(t) + N_m^i(t)$ will denote the total number of individuals of gender $i$. Then the following steps describe the life cycle to go from generation $t$ to $t+1$.

1. Encountering mates: Each wild-type female encounters a mate (male) with probability $1 - e^{-r_w N^M(t)}$, and each mutant female encounters a mate (male) with probability $1 - e^{-r_m N^M(t)}$. Upon encounter, mating is guaranteed.

2. Formation of mating pairs: Since the encounter is random, the male encountered is chosen randomly from the pool of all males. Therefore the probability that a female will mate with a wild-type or mutant male is $\dfrac{N_w^M(t)}{N^M(t)}$ or $\dfrac{N_m^M(t)}{N^M(t)}$ respectively. $N_{j_1 j_2}$ will denote the number of mating pairs with a female of type $j_1$ and male of type $j_2$.

3. Offspring production: The number of offspring of each mating pair is determined according to a Poisson distribution with mean $\lambda_w$ if the female is a wild type and $\lambda_m$ if the female is a mutant. The offspring inherit the genotype of either the mother or father with equal probability. Therefore the expected number of individuals of each type in the next generation (before mutation) is given by

$$\begin{aligned}\tilde{N}_w(t+1) &= \big(N_{ww}(t) + \frac{1}{2}N_{wm}(t)\big)\lambda_w + \frac{1}{2}N_{mw}(t)\lambda_m,\\ \tilde{N}_m(t+1) &= \big(N_{mm}(t) + \frac{1}{2}N_{mw}(t)\big)\lambda_m + \frac{1}{2}N_{wm}(t)\lambda_w.\end{aligned}$$

4. Mutation: Each wild-type offspring acquires the mutation with probability $\mu$. Therefore, the number of individuals in of each type post mutation in the next generation is given by

$$\begin{aligned}N_w(t+1) &\sim \text{Poisson}\big((1-\mu)\tilde{N}_w(t+1)\big),\\ N_m(t+1) &\sim \text{Poisson}\big(\mu\tilde{N}_w(t+1) + \tilde{N}_m(t+1)\big).\end{aligned}$$

5. Sex determination: Each individual is a female with probability $\alpha_{\text{fem}}$ and male otherwise:

$$\begin{aligned} N_j^F(t+1) &\sim \text{Binomial}(N_j(t+1), \alpha_{\text{fem}}), \\ N_j^M(t+1) &= N_j(t+1) - N_j^F(t+1). \end{aligned}$$

The life cycle is illustrated in Fig. 1 in the main text.

#### S1.2 Androdioecious population

Let $N_j^i(t)$ ($i = M, H$ , $j = w, m$) denote the number of individuals in generation $n$ of type $j$ (*w*ildtype or *m*utant) and gender $i$ (*M*ale or *H*ermaphoditic). The hermaphroditic individuals can reproduce sexually by mating with a male or a hermaphroditic individual, or undergo selfing. Further, $N_j(t) = N_j^M(t) + N_j^H(t)$ denotes the number of individuals of type $j$, $N^i(t) = N_w^i(t) + N_m^i(t)$ denotes the number of individuals of gender $i$ and $N(t) = N^H(t) + N^M(t)$ denotes the total number of individuals in the population. Then the following steps describe the life cycle to go from generation $t$ to $t + 1$.

1. Encountering mates: Each hermaphroditic wild-type individual encounters a mate (male or hermaphroditic) with probability $1-e^{-r_w(N(t)-1)}$. Each hermaphroditic mutant individual encounters a mate (male or hermaphroditic) with probability $1 - e^{-r_m(N(t)-1)}$. The mate pool size is $N - 1$ (instead of $N$) because an individual cannot mate with itself unless it selfs (see steps 2 and 3). Upon encounter, mating is guaranteed.

2. Formation of mating pairs: Since the encounter is random, the mating partner encountered is chosen randomly from the pool of all potential mates (i.e. both hermaphrodites and males). Since an individual cannot mate with itself unless it selfs, the probability that a hermaphrodite mates with a wild-type individual is $\dfrac{N_w(t) - 1}{N(t) - 1}$ or $\dfrac{N_w(t)}{N(t) - 1}$, depending on whether the hermaphroditic individual is a wild type or mutant respectively. Similarly, the probability that a hermaphrodite mates with a mutant individual is $\dfrac{N_m(t)}{N(t) - 1}$ or $\dfrac{N_m(t) - 1}{N(t) - 1}$ depending on whether the hermaphroditic individual is a wild type or mutant respectively. $N_{j_1 j_2}$ will denote the number of mating pairs with an individual of

type $j_1$ taking the role of the female and an individual of type $j_2$ taking the role of the male.

3. Selfing: Hermaphroditic individuals that did not obtain mates undergo selfing with probability $p_{\text{self}}$. The number of selfing individuals of a particular type $j$ is denoted by $N_j^{\text{self}}$.

4. Offspring production: The number of offspring of each mating pair is determined according to a Poisson distribution with mean $\lambda_j$ if they are outcrossing where $j$ is the type of the female of the mating pair. The offspring inherit the genotype of either of the parents with equal probability. Selfing individuals of type $j$ have a Poisson distributed number of offspring of type $j$ with mean $(1-\text{ID}_{\text{self}})\lambda_j$. Therefore the expected number of individuals of the two types in the next generation (before mutation) is given by

$$\begin{aligned}\tilde{N}_w(t+1) &= \big(N_{ww}(t) + \frac{1}{2}N_{wm}(t)\big)\lambda_w + \frac{1}{2}N_{mw}(t)\lambda_m + N_w^{\text{self}}(1-\text{ID}_{\text{self}})\lambda_w,\\ \tilde{N}_m(t+1) &= \big(N_{mm}(t) + \frac{1}{2}N_{mw}(t)\big)\lambda_m + \frac{1}{2}N_{wm}(t)\lambda_w + N_m^{\text{self}}(1-\text{ID}_{\text{self}})\lambda_m.\end{aligned}$$

5. Mutation: Each wild-type offspring acquires the mutation with probability $\mu$. Therefore, the number of individuals after mutation is given by

$$\begin{aligned}N_w(t+1) &\sim \text{Poisson}\big((1-\mu)\tilde{N}_w(t+1)\big),\\ N_w(t+1) &\sim \text{Poisson}\big(\mu\tilde{N}_w(t+1) + \tilde{N}_m(t+1)\big).\end{aligned}$$

6. Sex determination: Each individual is a hermaphrodite with probability $\alpha_{\text{her}}$ and male otherwise:

$$\begin{aligned}N_j^H(t+1) &\sim \text{Binomial}(N_j(t+1), \alpha_{\text{her}}),\\ N_j^M(t+1) &= N_j(t+1) - N_j^F(t+1).\end{aligned}$$

The life cycle is illustrated in Fig. S1(i).

#### S1.3 Hermaphroditic population

In hermaphroditic populations, all individuals are hermaphroditic, and the model is the same as that for the androdioecious population with $\alpha_{\mathrm{her}} = 1$. The life cycle is illustrated in Fig. S1(ii).

(a) **Life cycle in an androdioecious population**

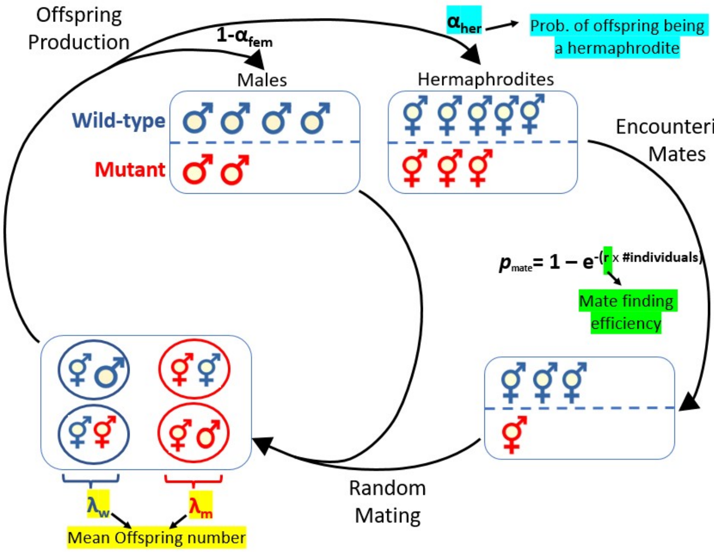


(b) **Life cycle in a hermaphroditic population**

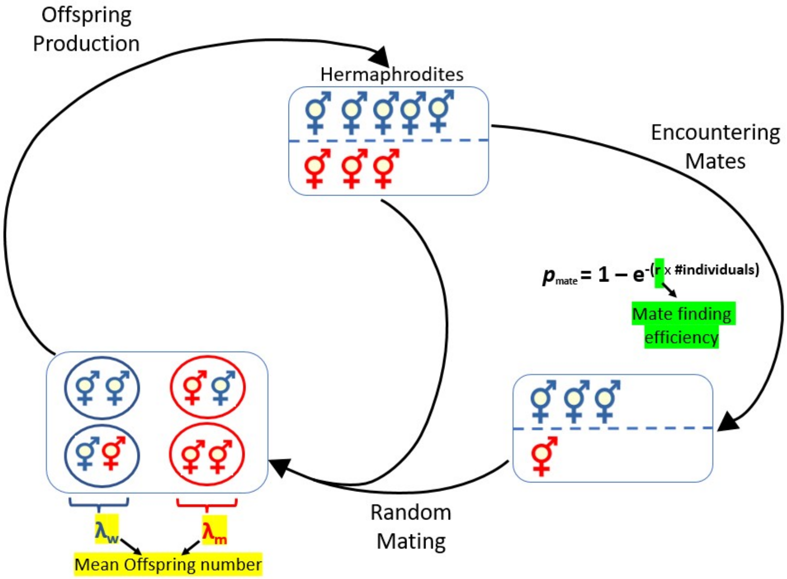


Figure S1: Diagrammatic representation of the stochastic life cycle to go from generation $t$ to $t+1$ for (a) androdioecious and (b) hermaphroditic populations (excluding selfing and mutation). The blue individuals have the wild-type phenotype and the red ones have the mutant phenotype. Note that hermaphroditic individuals can act as either a male or a female in a mating pair.

### Additional figures

Population dynamics without scaling the wild-type fecundity by the critical fecundity

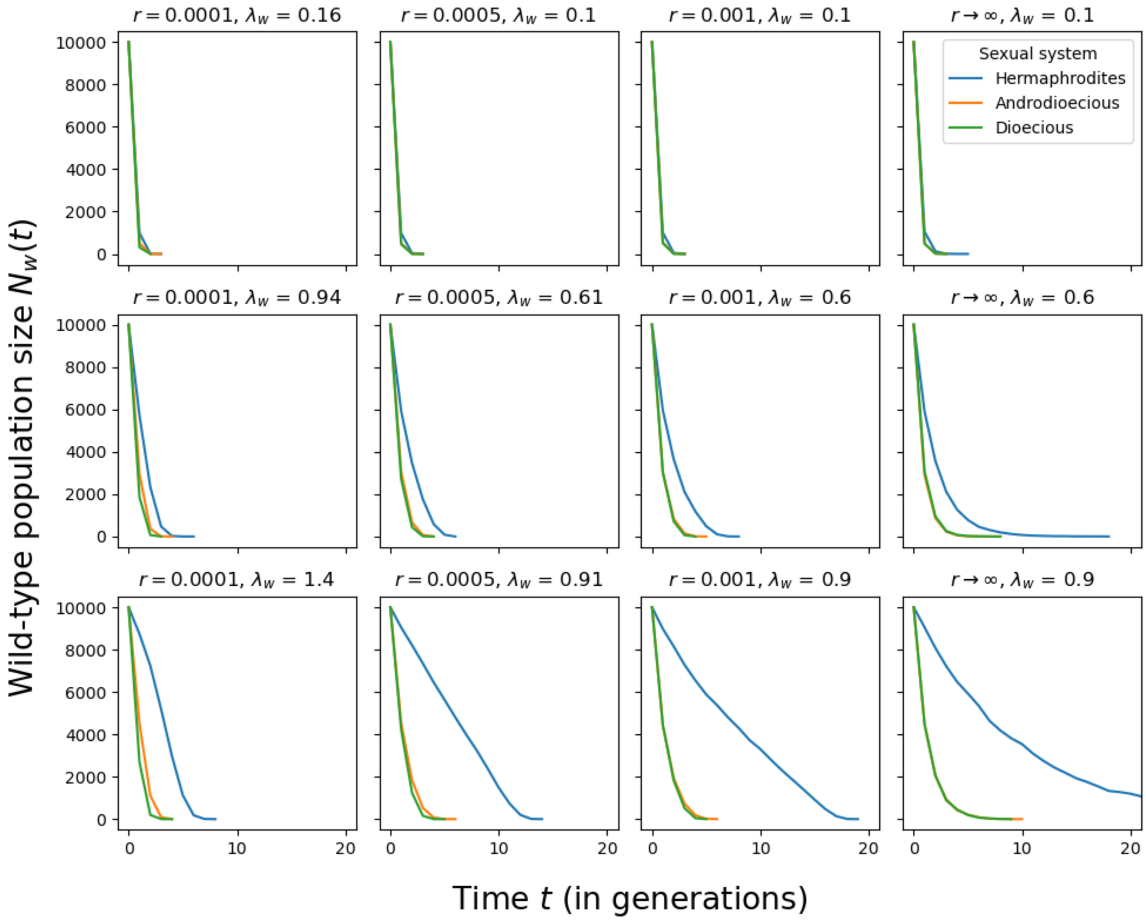


Figure S2: Population dynamics of the wild-type population (in the absence of mutations) across the three sexual systems. $\lambda_w$ is chosen such that the hermaphroditic system (blue line) has the population dynamics as in Fig. S3. The population dynamics, specifically the decay rate, differ substantially between the sexual systems.

Population dynamics with the wild-type fecundity scaled by the critical fecundity

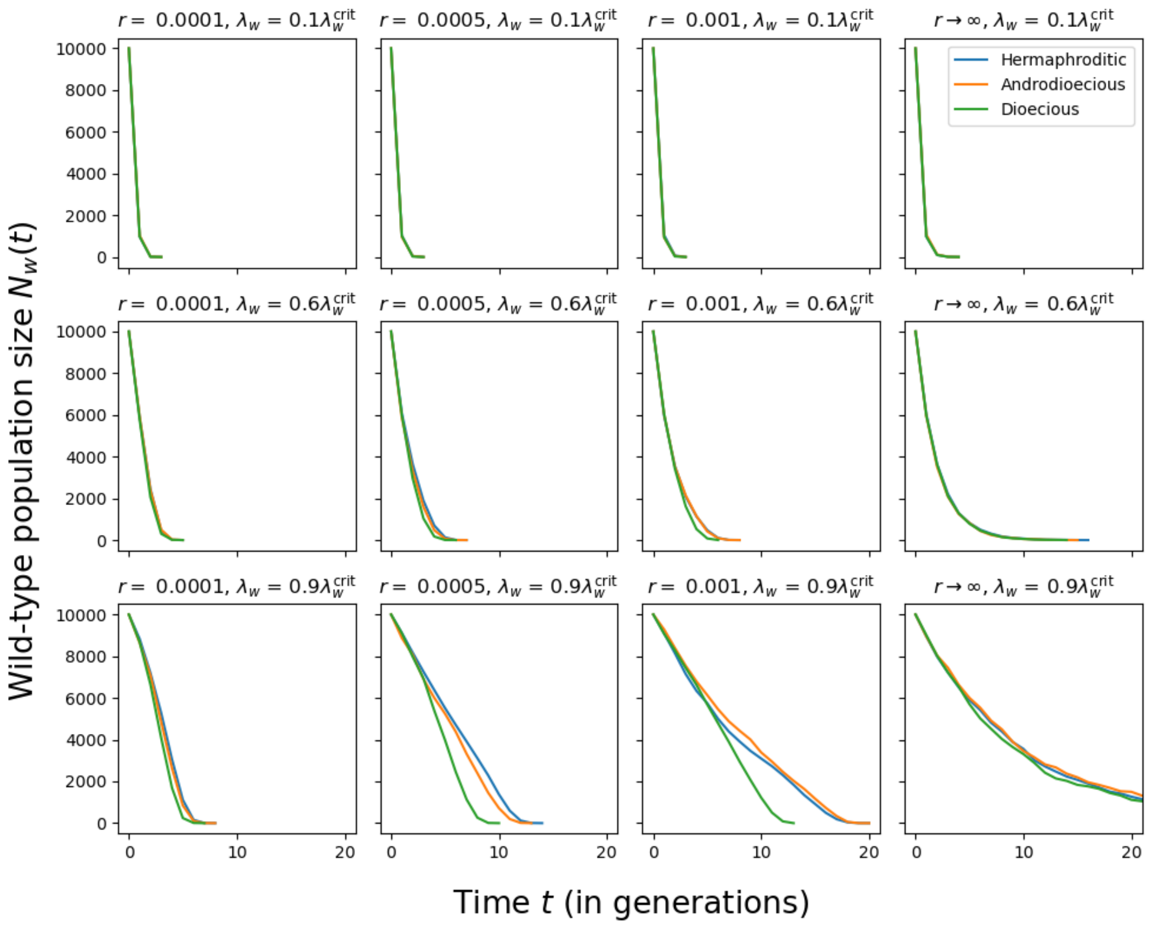


Figure S3: Population dynamics of the wild-type population (in the absence of mutations) across the three sexual systems when $\lambda_w/\lambda_w^{\text{crit}}$ is the same for all three populations. The decay rates of the populations are more similar than in Fig. S2. For a slow decay of the population size ($\lambda_w = 0.9\lambda_w^{\text{crit}}$), the three systems differ even with the scaling. We suspect that this is mostly because the difference in $\lambda_w^{\text{crit}}$ between androdioecious and dioecious populations is too small to be captured by our estimate, which only takes 2 decimal points into account; therefore the same value of $\lambda_w$ is used for both simulations.

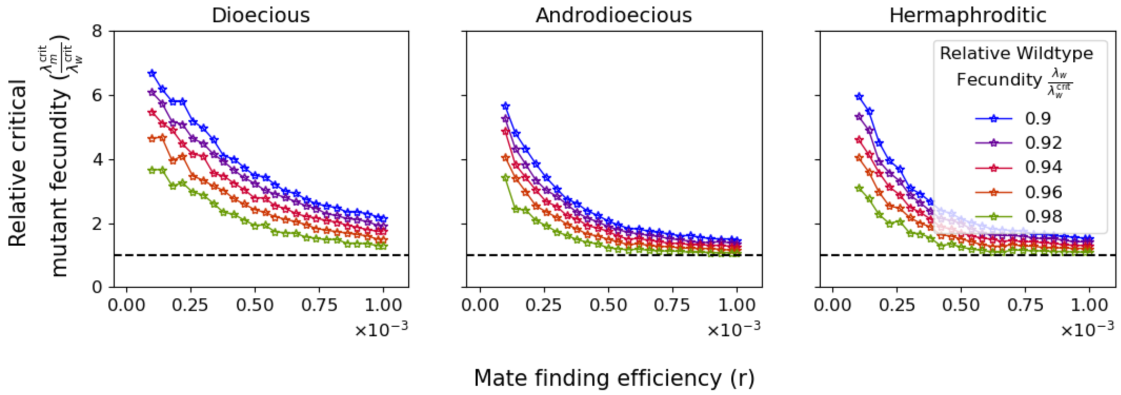


Figure S4: Critical mutant fecundity as in Fig. 2 (c) but scaled by the critical wildtype fecundity $\lambda_w^{\text{crit}}$.

### S2 Complementary analysis

#### S2.1 Expected change in the number of mutants from one generation to the next

In the following, we want to consider how the number of mutants changes from one generation to the next. For notational simplicity, we set $N(t) \equiv N$, $N_w(t) \equiv N_w$, $N_m(t) \equiv N_m$, $N_w(t+1) \equiv N'_w$, $N_m(t+1) \equiv N'_m$ (and equivalently for the numbers of males, females, and hermaphrodites). We consider two situations: (1) Proliferation of rare mutants when $N_m \ll N_w$ and mating between mutants can be ignored and (2) Spread of mutants in later generations when we cannot ignore mating of mutants with each other, where we assume that for a dioecious population $N_m^F = \alpha_{\text{fem}} N_m$ and $N_m^M = (1-\alpha_{\text{fem}})N_m$ and for an androdioecious population $N_m^F = \alpha_{\text{her}} N_m$ and $N_m^M = N_m$. In both situations (rare and frequent mutants), we set $N_w^F = \alpha_{\text{fem}} N_w$ and $N_w^M = (1-\alpha_{\text{fem}})N_w$ (dioecious population) and $N_w^F = \alpha_{\text{her}} N_w$ and $N_w^M = N_w$ (androdioecious population).

##### S2.1.1 Dioecious population

Following the model description in section S1.1, we obtain for the expected change in the number of mutants from one generation to the next:

$$\begin{aligned}\Delta N_m &= \mathbb{E}[N'_m] - N_m \\ &= p_{\text{mate}} \cdot \left( N_m^F \cdot \lambda_m \cdot \frac{N_m^M}{N_m^M + N_w^M} + N_m^F \cdot \frac{\lambda_m}{2} \cdot \frac{N_w^M}{N_m^M + N_w^M} + N_w^F \cdot \frac{\lambda_w}{2} \cdot \frac{N_m^M}{N_m^M + N_w^M} \right) - N_m\end{aligned} \tag{S2.1}$$

with $p_{\text{mate}} = 1 - e^{-rN^M}$.

**Proliferation of rare mutants.** When mutants are rare, $N_m \ll N_w$, we can ignore mating between mutants and furthermore approximate $N_w + N_m \approx N_w$. With this, we obtain:

$$\begin{aligned}\Delta N_m &\approx \left(1 - e^{-r(1-\alpha_{\text{fem}})N_w}\right) \cdot \left( \alpha_{\text{fem}} N_m \cdot \frac{\lambda_m}{2} + \alpha_{\text{fem}} N_w \cdot \frac{\lambda_w}{2} \cdot \frac{N_m}{N_w} \right) - N_m \\ &= \alpha_{\text{fem}} N_m \cdot \frac{\lambda_m + \lambda_w}{2} \cdot \left(1 - e^{-r(1-\alpha_{\text{fem}})N_w}\right) - N_m.\end{aligned} \tag{S2.2}$$

**Spread of frequent mutans.** When mutants are frequent (i.e. especially at later generations), we obtain

$$\begin{aligned}\Delta N_m &= \left(\alpha_{\text{fem}} N_m \lambda_m \frac{N_m}{N_m+N_w} + \alpha_{\text{fem}} N_m \frac{\lambda_m}{2}\frac{N_w}{N_m+N_w} + \alpha_{\text{fem}} N_w \frac{\lambda_w}{2}\frac{N_m}{N_m+N_w}\right) \\ &\quad \times \left(1-e^{-r(1-\alpha_{\text{fem}})(N_w+N_m)}\right) - N_m \\ &= \alpha_{\text{fem}} N_m \frac{\lambda_m N_m + \frac{\lambda_m+\lambda_w}{2} N_w}{N_m+N_w} \cdot \left(1-e^{-r(1-\alpha_{\text{fem}})(N_w+N_m)}\right) - N_m.\end{aligned} \tag{S2.3}$$

##### S2.1.2 Androdioecious population

Following the model description in section S1.2, we obtain for the expected change in the number of mutants from one generation to the next:

$$\begin{aligned}&\Delta N_m = \mathbb{E}[N_m'] - N_m \\ &= p_{\text{mate}} \cdot \left(N_m^H \cdot \lambda_m \cdot \frac{N_m-1}{N_m+N_w-1} + N_m^H \cdot \frac{\lambda_m}{2} \cdot \frac{N_w}{N_m+N_w-1} + N_w^H \cdot \frac{\lambda_w}{2} \cdot \frac{N_m}{N_m+N_w-1}\right) - N_m\end{aligned} \tag{S2.4}$$

with $p_{\text{mate}} = 1 - e^{-r(N-1)}$.

**Proliferation of rare mutants.** When mutants are rare, $N_m \ll N_w$, we can ignore mating between mutants and furthermore approximate $N_w + N_m - 1 \approx N_w$. With this, we obtain:

$$\begin{aligned}\Delta N_m &\approx \left(1-e^{-rN_w}\right) \cdot \left(\alpha_{\text{her}} N_m \cdot \frac{\lambda_m}{2} + \alpha_{\text{her}} N_w \cdot \frac{\lambda_w}{2} \cdot \frac{N_m}{N_w}\right) - N_m \\ &= \alpha_{\text{her}} N_m \cdot \frac{\lambda_m+\lambda_w}{2} \cdot \left(1-e^{-rN_w}\right) - N_m.\end{aligned} \tag{S2.5}$$

**Spread of frequent mutants.** For spread of mutants in later generations when $N_m \gg 1$, we obtain

$$\begin{aligned}\Delta N_m &= \left(\alpha_{\text{her}} N_m \lambda_m \frac{N_m - 1}{N_m + N_w - 1} + \alpha_{\text{her}} N_m \frac{\lambda_m}{2} \frac{N_w}{N_m + N_w - 1} + \alpha_{\text{her}} N_w \frac{\lambda_w}{2} \frac{N_m}{N_m + N_w - 1}\right) \\ &\quad \times \left(1 - e^{-r(N_w + N_m - 1)}\right) - N_m \\ &= \alpha_{\text{her}} N_m \frac{\lambda_m (N_m - 1) + \frac{\lambda_m + \lambda_w}{2} N_w}{N_m + N_w - 1} \cdot \left(1 - e^{-r(N_w + N_m - 1)}\right) - N_m. \end{aligned} \tag{S2.6}$$

#### S2.2 The change of relative frequency from one generation to the next

To complement the picture, we here derive the change in the relative frequency of the mutant allele from one generation to the next. We use the notation $p_m \equiv p_m(t)$ and $p'_m \equiv p_m(t+1)$. From Eq. (S2.3), we know that

$$N'_m = \alpha_{\text{fem}} N_m \frac{\lambda_m N_m + \frac{\lambda_m + \lambda_w}{2} N_w}{N_m + N_w} \cdot p_{\text{mate}}. \tag{S2.7}$$

Analogously, we have

$$N'_w = \alpha_{\text{fem}} N_w \frac{\lambda_w N_w + \frac{\lambda_m + \lambda_w}{2} N_m}{N_m + N_w} \cdot p_{\text{mate}}. \tag{S2.8}$$

With this and setting $\lambda_m = (1+s)\lambda_w$, we obtain

$$\begin{aligned} p'_m &= \frac{N'_m}{N'_w + N'_m} \\ &= \frac{p_m((1+s)p_m + (1+\frac{s}{2})(1-p_m))}{p_m(1+\frac{s}{2}+\frac{s}{2}p_m) + (1-p_m)(1+\frac{s}{2}p_m)} \\ &= \frac{p_m(1+\frac{s}{2}+\frac{s}{2}p_m)}{1} \\ &= \frac{p_m \cdot \left(1+\frac{s}{2}+\frac{s}{2}p_m\right)}{1+sp_m}. \end{aligned} \tag{S2.9}$$

Further:

$$\Delta p_m = p'_m - p_m = \frac{p_m + \frac{s}{2}p_m(1+p_m) - p_m - sp_m^2}{1+sp_m} = \frac{\frac{s}{2}p_m(1-p_m)}{1+sp_m}, \tag{S2.10}$$

which corresponds to the frequency change of an allele at a single diploid locus with selection coefficient $s$ and dominance coefficient $h = \frac{1}{2}$ (Equation 1.20 of Rice (2004)).

Similarly, for an androdioecious population we have:

$$N'_m = \alpha_{\text{her}} N_m \frac{\lambda_m (N_m - 1) + \frac{\lambda_m + \lambda_w}{2} N_w}{N_m + N_w - 1} \cdot p_{\text{mate}}, \tag{S2.11a}$$

$$N'_w = \alpha_{\text{her}} N_w \frac{\lambda_w (N_w - 1) + \frac{\lambda_m + \lambda_w}{2} N_m}{N_m + N_w - 1} \cdot p_{\text{mate}}. \tag{S2.11b}$$

From this:

$$\begin{aligned}
p'_m &= \frac{N'_m}{N'_w + N'_m} \\
&= \frac{(1+s)N_m(N_m - 1) + (1 + \frac{s}{2})N_m N_w}{(1+s)N_m(N_m-1) + N_w(N_w - 1) + (2+s)N_w N_m} \\
&= \frac{(1+s)p_m^2 + (1+\frac{s}{2})p_m(1-p_m) - (1+s)p_m \frac{1}{N_m+N_w}}{(1+s)p_m^2 + (1-p_m)^2 + (2+s)p_m(1-p_m) - (1+s)p_m\frac{1}{N_m+N_w} - (1-p_m)\frac{1}{N_m+N_w}} \\
&= \frac{p_m(1 + \frac{s}{2}(1+p_m)) - (1+s)p_m\frac{1}{N_m+N_w}}{1 + sp_m - (1+sp_m)\frac{1}{N_m+N_w}}
\end{aligned} \tag{S2.12}$$

and

$$\begin{aligned}
\Delta p_m &= p'_m - p_m \\
&= \frac{p_m(1-p_m)\frac{s}{2} - p_m s(1-p_m)\frac{1}{N_m+N_w}}{1 + sp_m - (1+sp_m)\frac{1}{N_m+N_w}} \\
&= \frac{p_m(1-p_m)\frac{s}{2}}{1+sp_m} \times \frac{1 - \frac{2}{N_m+N_w}}{1 - \frac{1}{N_m+N_w}} \\
&= \frac{p_m(1-p_m)\frac{s}{2}}{1+sp_m} \cdot \left(1 - \frac{1}{N_m + N_w - 1}\right) \\
&\approx \frac{\frac{s}{2}p_m(1-p_m)}{1+sp_m}.
\end{aligned} \tag{S2.13}$$

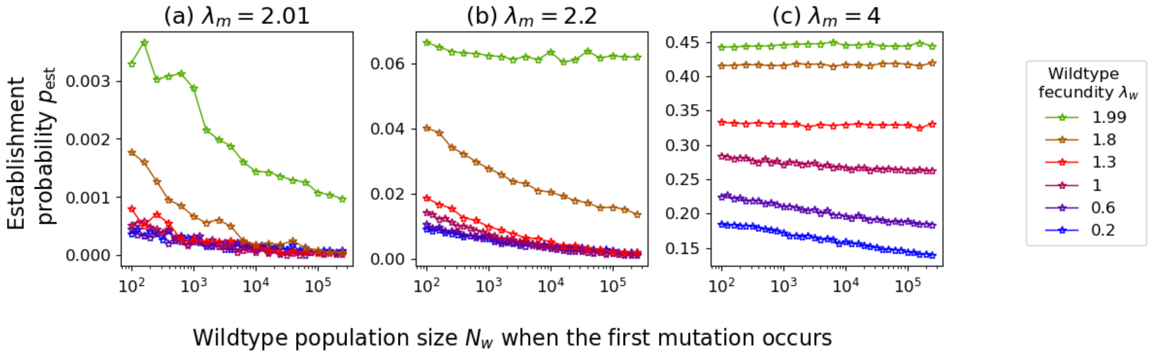


Figure S5: The establishment probability of a single mutant in a dioecious population with assured mating as a function of the wild-type population size at the time of appearance of the mutation.

#### S2.3 The role of the wild-type population size in the spread of mutants

In this section, we want to determine how $\Delta N_m$ depends on the wild-type population size $N_w$. On the one hand, a large wild-type population size increases the probability of mating. On the other hand, mutants are less likely to mate with each other (provided mating occurs) if the wild-type population size is large. From Eq. (S2.2) and (S2.5), we immediately see that in the presence of mate limitation, proliferation of rare mutants is more likely if the wild-type population size is large in both dioecious and androdioecious populations; in the absence of mate limitation ($r \to \infty$), it is independent of $N_w$.

We next turn to the spread of mutants when mating between mutants is not negligible (Eq. S2.3 and Eq. S2.6). To perform the calculation simultaneously for dioecious and androdioecious populations, we introduce the parameters $\alpha^F$, $\alpha^M$, and $\delta$. For a dioecious population, we have $\alpha^F = \alpha_{\text{fem}}$, $\alpha^M = 1 - \alpha_{\text{fem}}$, and $\delta = 0$. For an androdioecious population, we have $\alpha^F = \alpha_{\text{her}}$, $\alpha^M = 1$, and $\delta = 1$. We furthermore choose the notation $N_w \equiv x$ such that we can express $\mathbb{E}[N_m(t+1)]$ as a function of the variable $x$:

$$g(x) = f(x) \cdot (1 - e^{-r\alpha^M(N_m + x - \delta)}) \tag{S2.14}$$

with

$$
\begin{aligned}
f(x) &= \alpha^F N_m \frac{\lambda_m(N_m-\delta)+\frac{\lambda_m+\lambda_w}{2}x}{N_m+x-\delta}. \\
&= \alpha^F N_m \left(\lambda_m + \frac{\lambda_w-\lambda_m}{2}\frac{x}{N_m+x-\delta}\right)
\end{aligned}
\tag{S2.15}
$$

We will look at the limit $r\to\infty$ and at very small $r$.

In the limit $r\to\infty$, we have $p_{\text{mate}}=1$. We thus now look at the function $f(x)$. Taking the derivative yields

$$
\begin{aligned}
f'(x) &= \alpha^F N_m \frac{\lambda_w-\lambda_m}{2}\frac{N_m+x-\delta-x}{(N_m+x-\delta)^2} \\
&= \alpha^F N_m \frac{\lambda_w-\lambda_m}{2}\frac{N_m-\delta}{(N_m+x-\delta)^2} < 0.
\end{aligned}
\tag{S2.16}
$$

where the inequality at the end holds since $\lambda_w<\lambda_m$. I.e., $\mathbb{E}[N_m(t+1)]$ decreases with $N_w$. This means that $\Delta N_m$ is smaller for larger $N_w$.

For finite $r$, we need to look at the derivative of $g(x)$. We obtain

$$
\begin{aligned}
g'(x) &= \alpha^F N_m \frac{\lambda_w-\lambda_m}{2}\frac{N_m-\delta}{(N_m+x-\delta)^2}\left(1-e^{-r\alpha^M(N_m+x-\delta)}\right) \\
&\quad + \alpha^F N_m\left(\lambda_m+\frac{\lambda_w-\lambda_m}{2}\frac{x}{N_m+x-\delta}\right) r\alpha^M e^{-r\alpha^M(N_m+x-\delta)} \\
&\geq \alpha^F N_m \frac{\lambda_w-\lambda_m}{2}\frac{N_m-\delta}{N_m+x-\delta} r\alpha^M + \\
&\quad + \alpha^F N_m\left(\lambda_m+\frac{\lambda_w-\lambda_m}{2}\frac{x}{N_m+x-\delta}\right) r\alpha^M\left(1-r\alpha^M(N_m+x-\delta)\right) \\
&= r\alpha^F\alpha^M N_m \times \left(\frac{\lambda_w-\lambda_m}{2}\frac{N_m-\delta}{N_m+x-\delta}+\lambda_m+\frac{\lambda_w-\lambda_m}{2}\frac{x}{N_m+x-\delta}\right. \\
&\quad \left. - \big[\lambda_m+\frac{\lambda_w-\lambda_m}{2}\frac{x}{N_m+x-\delta}\big] r\alpha^M(N_m+x-\delta)\right) \\
&= r\alpha^F\alpha^M N_m \times \left(\frac{\lambda_w+\lambda_m}{2} - \big[\lambda_m+\frac{\lambda_w-\lambda_m}{2}\frac{x}{N_m+x-\delta}\big] r\alpha^M(N_m+x-\delta)\right) \\
&= r\alpha^F\alpha^M N_m \times \left(\frac{\lambda_w+\lambda_m}{2} - \big[\lambda_m(N_m+\frac{x}{2}-\delta)+\lambda_w\frac{x}{2}\big] r\alpha^M\right),
\end{aligned}
\tag{S2.17}
$$

where the inequality from step 1 to 2 holds since $e^{-ay}\geq 1-ay$ for $a>0$ and the first term is negative while the second term is positive. The second term in the bracket in the last line of Eq. (S2.17) is always smaller than the first one if $r$ is sufficiently small, which shows that for small enough $r$, $g'(x) > 0$. This means that for sufficiently small $r$, $\mathbb{E}[N_m(t+1)]$ and thus $\Delta N_m$ increases with $N_w(t)$.

If the mutation appears in an otherwise wild-type population, it will either get lost again while rare or establish a long-term lineage of offspring that rescues the population. Considering the dependence of $\Delta N_m$ on $N_w$ provides us with insights about the establishment probability $p_{\text{est}}$ of the rescue mutation. For small mate-finding efficiencies $r$, we have seen that $\Delta N_m$ increases with an increasing wild-type population size, and we expect the same for $p_{\text{est}}$, i.e. mutants that appear when the population size is large, have a larger establishment probability. This is intuitive since the wild-type population size has a strong effect on their mate-finding probability. We next turn to assured mating ($r \to \infty$). Although mutants will mate independent of the wild-type population size (as long as there is at least one male to mate with), we have seen that the wild-type population size nevertheless can affect the spread of mutants, albeit more subtly: when the relative frequency of mutants is sufficiently high that mating between them is non-negligible, $\Delta N_m$ decreases with increasing $N_w$ since a large wild-type population size makes it less likely for mutants to mate with each other. However, when the mutant fecundity $\lambda_m$ is small, it is necessary for mutants to mate with each other for successful establishment (otherwise, only half of the offspring of a mutant are mutants, which are too few to allow for their spread). Since mating between mutants is more likely when the wild-type population size is small, the establishment probability decreases with increasing wild-type population size at the time of mutant appearance (Fig. S5a and b) in line with the decrease in $\Delta N_m$ with $N_w$. When $\lambda_m$ is large, mutants do not need to mate with each other for successful establishment, and the establishment probability is independent of the wild-type population size (Fig. S5c). This is consistent with the independence of $\Delta N_m$ from $N_w$ for rare mutants.

#### S2.4 The effect of the mutant fecundity on the comparison between $P_{\text{res}}^{(\text{SGV})}$ and $P_{\text{res}}^{(\text{DNM})}$

In Fig. 4 of the main text, we observe that the wild-type fecundity $\hat{\lambda}_w$ for which $P_{\text{res}}^{(\text{SGV})} = P_{\text{res}}^{(\text{DNM})}$ depends on the mutant fecundity $\lambda_m$: In the limit $r \to \infty$, this wild-type fecundity $\hat{\lambda}_w$ increases with increasing $\lambda_m$, while for small $r$, it decreases. We here provide an approximate argument why these shifts occur based on the rationale outlined in the section "Preliminary considerations", making use of Eq. 3.

If the establishment probability $p_{\text{est}}(t)$ were independent of time, i.e. $p_{\text{est}}(t) = \text{const.}$, the wild-type fecundity $\hat{\lambda}_w$ would be entirely determined by $N_m(0) = \sum_{t=0}^{\infty} N_{\text{dnm}}(t)$. However, as we have already seen in section S2.3 and as is also intuitively expected, the establishment probability depends on the wild-type population size and thus on time (since $N_w$ decreases over time). Further, if the establishment probability decreases (resp. increases) with time then $\hat{\lambda}_w$ will be higher (resp. lower) compared to the case with a constant establishment probability. For the arguments to follow we will use the dependence of $p_{\text{est}}$ on $N_w$ (see section S2.3) as a proxy for the dependence of the establishment probability $p_{\text{est}}$ on time; decreasing $N_w$ corresponds to increasing time $t$. In section S2.3, we found that $p_{\text{est}}$ increases with decreasing $N_w$ for $r \to \infty$ (for small $\lambda_m$). This means that the establishment probability increases with time, which implies that $\hat{\lambda}_w$ is lower than it would be for a constant establishment probability. By contrast, we argued based on $\Delta N_m$, that $p_{\text{est}}$ increases with $N_w$ for small $r$ such that late-occurring mutations have a lower establishment probability and $\hat{\lambda}_w$ is higher than it would be for a constant establishment probability.

We now argue that the effect is stronger for small than for large mutant fecundities $\lambda_m$. We have already seen this for assured mating, where the establishment probability strongly depends on $N_w$ for small $\lambda_m$ but is independent of $N_w$ for large $\lambda_m$ (section S2.3 and Fig. S5). This explains why in Fig. 4a, $\hat{\lambda}$ is mostly independent of $\lambda_m$ within the range of large $\lambda_m$ and only shifts to the left for small $\lambda_m$. We next turn to small $r$. The number of offspring that a mutant produces depends both on its probability to mate $p_{\text{mate}}$ (which strongly depends on $N_w$) and on $\lambda_m$. The smaller $\lambda_m$, the more important it is that $p_{\text{mate}}$ is high. Put differently, if the first mutant only produces few offspring (low $\lambda_m$), it is important for establishment that these offspring manage to mate themselves. However, if it produces many offspring (large $\lambda_m$), it is less important that each of them finds a mate. Therefore, a large wild-type population is more important for mutant establishment for small than for large $\lambda_m$. Consequently, $\hat{\lambda}_w$ increases as $\lambda_m$ decreases in Fig. 4b and c.

While Fig. 4 shows results for $N_m(0) = 1$, Fig. S6 shows results for $N_m(0) = 10$, finding the same trend.

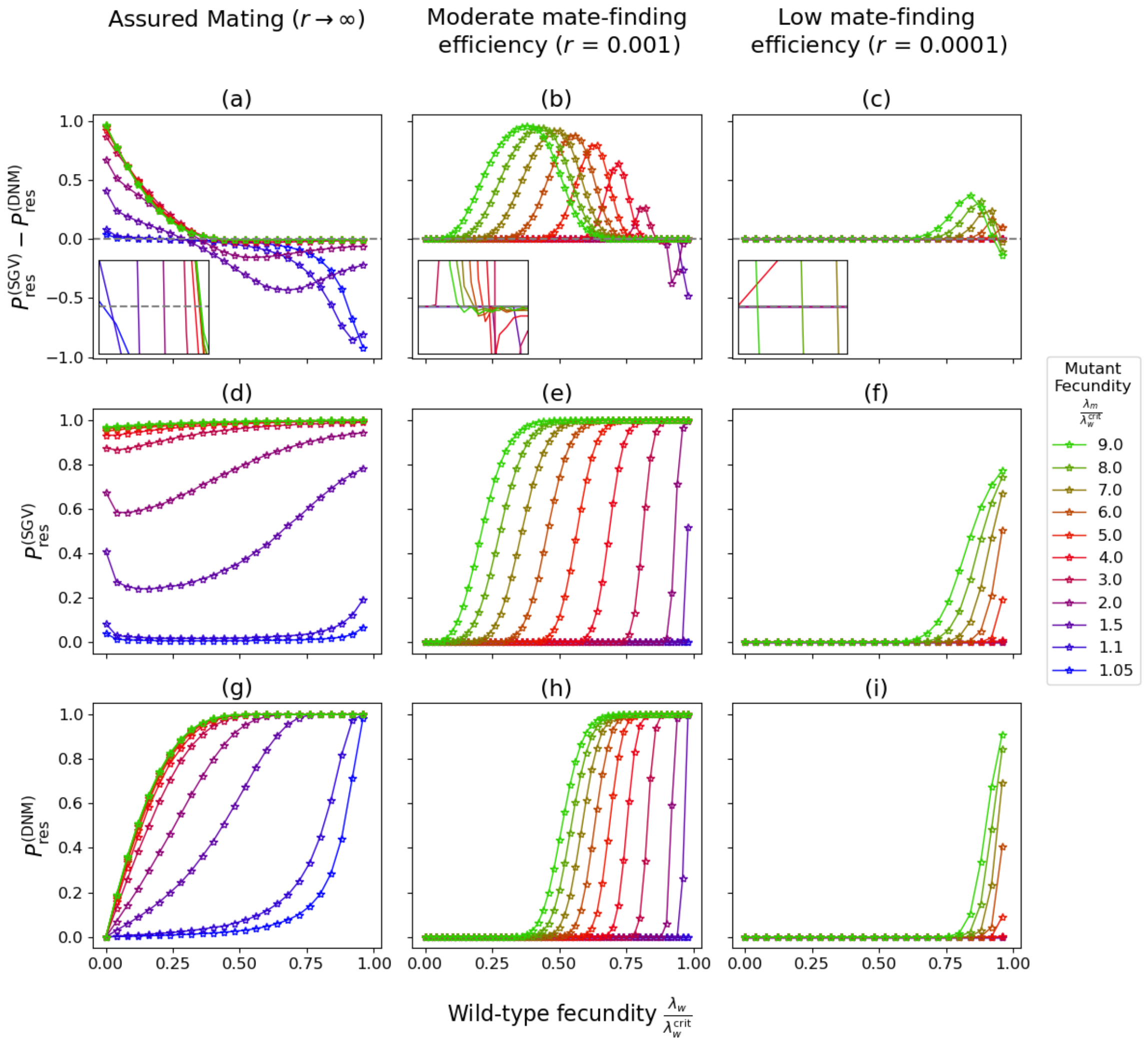


Figure S6: Comparison of the probabilities of rescue from the standing genetic variation $P_{\mathrm{res}}^{(\mathrm{SGV})}$) and from *de novo* mutations $P_{\mathrm{res}}^{(\mathrm{DNM})}$ in a dioecious population. The figure corresponds to Fig. 4 in the main text with the difference that we set $N_m(0) = 10$ in the present figure.

#### S2.5 The dependence of the optimal sex ratio on the mate-finding efficiency

We return to the approximation for the probability of rescue from *de novo* mutations given in Eq. (3):

$$P_{\mathrm{res}}^{(\mathrm{DNM})} = 1 - \exp\Big(-\sum_{t=0}^{\infty} N_{\mathrm{dnm}}(t) \cdot p_{\mathrm{est}}(t+1)\Big).$$

We want to know for which sex ratio, the exponent and thus the probability of rescue is largest.

###### S2.5.1 Dioecious population

The (expected) number of new mutations from generation $t$ to $t+1$, $N_{\text{dnm}}(t)$, is given by

$$N_{\text{dnm}}(t) = \mu \alpha_{\text{fem}} N_w(t) \lambda_w p_{\text{mate}}(t) \tag{S2.18}$$

with $p_{\text{mate}} = 1 - e^{-r(1-\alpha_{\text{fem}})N_w}$. As a proxy for the dependence of the establishment probability $p_{\text{est}}$ on $\alpha_{\text{fem}}$, we consider the dependence of $\Delta N_m$ on $\alpha_{\text{fem}}$ given by Eqs. (S2.2) and (S2.3).

Following the model description in section S1.1, the wild-type population size changes in the absence of mutants according to

$$\mathbb{E}[N_w(t+1)] = \alpha_{\text{fem}} N_w(t) \lambda_w \left(1 - e^{-r(1-\alpha_{\text{fem}})N_w}\right), \tag{S2.19}$$

which we will need in the following argument (see Eq. 4a in the main text).

**Assured mating.** We first consider $r \to \infty$, in which case $p_{\text{mate}} = 1$. The wild-type population size then decays slowest for $\alpha_{\text{fem}} \approx 1$ (see Eq. S2.19). (The proportion of female offspring $\alpha_{\text{fem}}$ needs to be a little lower than one, since some male is needed for mating.) Likewise, the rate of new mutations is maximal for $\alpha_{\text{fem}} \approx 1$ (see S2.18). From Eqs. (S2.2), we immediately see that $\Delta N_m$ for the proliferation of rare mutants also increases with $\alpha_{\text{fem}}$. The picture is more complicated for the spread of mutants when mating between them cannot be ignored: The direct dependence of $\Delta N_m$ of $\alpha_{\text{fem}}$ in Eq. (S2.3) is positive. However, we know from section S2.3 that $\Delta N_m$ decreases with $N_w$. This suggests that spread of mutants for $N_m \gg 1$ might be highest if the proportion is somewhat lower than one. This is generally expected to be a minor effect compared to the generation of new mutants and the proliferation of rare mutants but it slightly affects the optimal sex ratio when the wild-type population decays fast and the mutant fecundity is low (Fig. S7a). We overall conclude that, apart from this exception, $\alpha_{\text{fem}} \approx 1$ (slightly below one) is optimal in the limit $r \to \infty$.

**Low mate-finding efficiency.** For low $r$, we can approximate $p_{\text{mate}} \approx r(1-\alpha_{\text{fem}})(N_w + N_m)$ (by Taylor expansion in $r$). The wild-type population size then decays most slowly for $\alpha_{\text{fem}} \approx 0.5$. The number of new mutations $N_{\text{dnm}}(t)$ and $\Delta N_m + N_m$

for proliferation of rare mutants are both proportional to $\alpha_{\text{fem}}(1 - \alpha_{\text{fem}})N_w(t)$ and are thus largest for $\alpha_{\text{fem}} = 0.5$. The considerations for $\Delta N_m$ for spread of mutants in later generations is again more complicated. The term $\Delta N_m + N_m$ is proportional to $\alpha_{\text{fem}}(1 - \alpha_{\text{fem}})$, and we know from section S2.3 that it increases in $N_w$. Spread of mutants for $N_m \gg 1$ is thus highest for $\alpha_{\text{fem}} = 0.5$. We can therefore conclude that an equal sex ratio maximizes the probability of rescue for low mate-finding efficiencies $r$.

##### S2.5.2 Androdioecious population

The (expected) number of new mutations from generation $t$ to $t+1$, $N_{\text{dnm}}(t)$, is given by

$$N_{\text{dnm}}(t) = \mu \alpha_{\text{her}} N_w(t) \lambda_w p_{\text{mate}}(t) \tag{S2.20}$$

with $p_{\text{mate}} = 1 - e^{-r(N_w - 1)}$. As a proxy for the dependence of the establishment probability $p_{\text{est}}$ on $\alpha_{\text{het}}$, we consider the dependence of $\Delta N_m$ on $\alpha_{\text{fem}}$ given by Eqs. (S2.5) and (S2.6).

Following the model description in section S1.2, the wild-type population size changes in the absence of mutants according to

$$\mathbb{E}[N_w(t+1)] = \alpha_{\text{her}} N_w(t) \lambda_w \left(1 - e^{-r(N_w - 1)}\right), \tag{S2.21}$$

which we will need in the following argument (see Eq. 5a in the main text).

We immediately see that the wild-type population decays slowest and $N_{\text{dnm}}$ is highest for $\alpha_{\text{her}} = 1$, irrespective of the mate-finding efficiency. We next look at the proliferation of rare mutants (Eq. S2.5). Since $N_w$ increases with $\alpha_{\text{her}}$, irrespective of $r$, $\Delta N_m$ for the proliferation of rare mutants is highest for $\alpha_{\text{her}} = 1$. $\Delta N_m$ for the spread of mutants in later generations (Eq. S2.6) is directly proportional to $\alpha_{\text{her}}$ but also depends on $N_w$. From section S2.3, we know that, when mating between mutant individuals cannot be ignored, $\Delta N_m + N_m$ decreases with $N_w$ for assured mating and increases with $N_w$ for low $r$. This means that for low $r$, $\Delta N_m$ is highest for $\alpha_{\text{her}} = 1$, while for large $r$, slightly lower $\alpha_{\text{her}}$ could be better. However, as for the dioecious population, the latter is expected to be a minor effect. Indeed, we have never observed it (see e.g. Fig. S7b), but we can of course not exclude that it exists in some parameter regime that we did not explore. We can overall conclude that rescue is highest for $\alpha_{\text{her}} = 1$, i.e. for a hermaphroditic population.

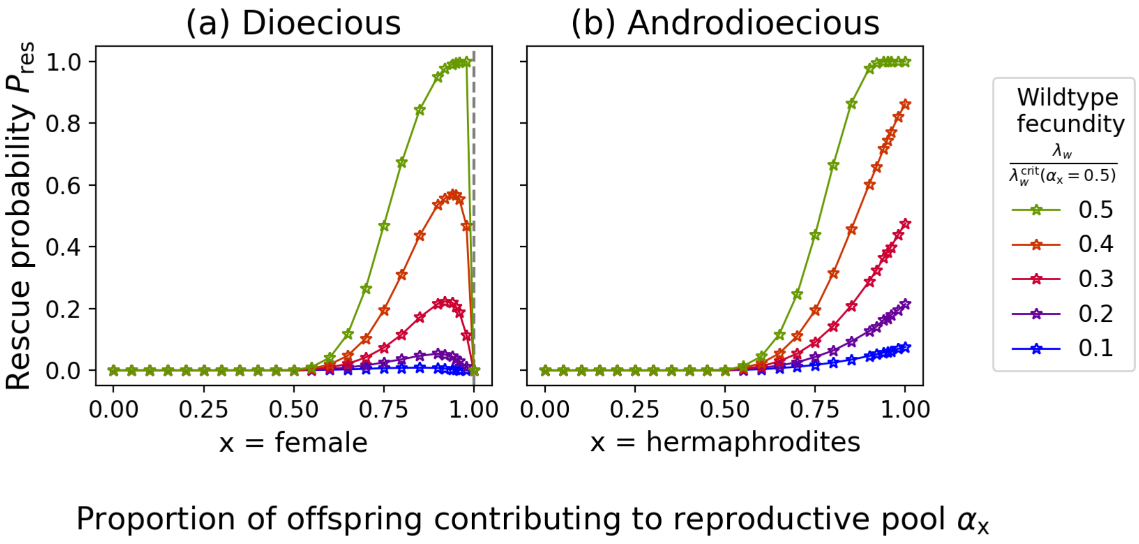


Figure S7: Probability of rescue as a function of proportion of female or proportion of hermaphrodites when mates are assured for dioecious and androdioecious populations. $\lambda_m = 1.02\lambda_w^{\text{crit}}, \mu = 10^{-4}, N_0 = 10^4$
